# PhageLysData: an evidence-aware and AI-ready dataset of phage lytic enzymes and depolymerases

**DOI:** 10.64898/2026.08.24.746620

**Authors:** David Medina-Ortiz, Álvaro Olivera-Nappa, María Elena Lienqueo, Rafael Opazo, Jaime Romero

**Affiliations:** Departamento de Ingeniería En Computación, Universidad de Magallanes, Avenida Bulnes 01855, 6210427, Punta Arenas, Chile; Centro de Biotecnología y Bioingeniería (CeBiB), Departamento de Ingeniería Química, Biotecnología y Materiales, Universidad de Chile, Av. Beauchef 851, 8370458, Santiago, Chile; Laboratorio de Biotecnología de Alimentos, Instituto de Nutrición y Tecnología de los Alimentos (INTA), Universidad de Chile, El Líbano 5524, Santiago 7830489, Chile

**Keywords:** Bacteriophage lytic enzymes, Phage depolymerases, Evidence-aware data integration, Exact-sequence integration, Data provenance, Protein language models, Functional annotation, FAIR data

## Abstract

Bacteriophage lytic enzymes and depolymerases are relevant to phage biology, antimicrobial development, and protein engineering, but their sequence and annotation data remain dispersed across general databases, specialized resources, genome-centred collections, and prediction-oriented datasets. We present PhageLysData, an evidence-aware and AI-ready resource constructed through reproducible multisource integration, provenance tracking, and exact-sequence consolidation. The release integrates 807,366 source observations from seven primary resources into 759,105 unique exact-sequence entities, comprising an evidence-supported Core of 11,867 entities, a Prediction Extension of 745,092 prediction-only candidates, and 2,146 Context entities retained for provenance and reference. This architecture preserves broad sequence-space coverage while maintaining a clear distinction between non-predictive and prediction-derived support. Core entities are enriched with harmonized biological annotations, physicochemical properties, independent InterProScan-derived functional annotations, mapped PDB and AlphaFold DB structural assets, and reusable numerical representations. For 11,259 eligible Core sequences, PhageLysData provides embeddings from 11 protein language models together with one-hot encoding under a common representation contract. Release-facing examples demonstrate latent-space exploration, unsupervised clustering, supervised classification, and evidence-aware candidate retrieval without defining a universal predictive benchmark. PhageLysData provides a traceable, versioned, and computationally accessible foundation for protein retrieval, comparative analysis, task-specific dataset construction, and machine-learning applications involving phage lytic enzymes and depolymerases.

## 1 Introduction

Bacteriophages encode enzymes that modify or degrade bacterial cell-envelope and extracellular structures during adsorption, genome entry, and progeny release (Dicks and Vermeulen, 2024; Fernandes and São-José, 2018). Endolysins hydrolyze peptidoglycan to promote host-cell lysis, virion-associated lytic enzymes mediate localized cell-wall degradation during infection, and phage depolymerases target capsular polysaccharides, extracellular polysaccharides, lipopolysaccharides, and related surface barriers (Grishin et al., 2020; Zhydzetski et al., 2024; Guliy and Evstigneeva, 2025; Cheetham et al., 2024; Topka-Bielecka et al., 2021). These proteins span diverse catalytic mechanisms, substrates, domain architectures, and biological roles, making them relevant to phage biology, antimicrobial development, enzyme mining, and protein engineering (Knecht et al., 2020; Lee et al., 2023). Their diversity has also motivated computational approaches for candidate discovery and comparative sequence and structural analysis (Li et al., 2020; Boulay et al., 2026).

The expansion of phage genome and protein collections has substantially increased the searchable sequence space for these enzymes (Cook et al., 2021). Relevant information is nevertheless distributed across general protein repositories, specialized phage-enzyme resources, genome-centred collections, and prediction-oriented datasets (uni, 2025; Criel et al., 2021; Magill and Skvortsov, 2023; Russell and Hatfull, 2017). Functional nomenclature, catalytic annotations, structural terminology, taxonomic constraints, and source-specific prediction systems retrieve overlapping but non-equivalent candidate sets, such that no single database, annotation field, or search term provides a complete and consistently defined collection (Latka et al., 2017; Alrafaie and Stafford, 2023).

Multisource integration introduces a second challenge because database occurrence is not equivalent to independent biological evidence. Identical protein sequences can occur under different accessions, genomes, or source-specific identifiers, while specialized resources may reuse annotations, predictions, publications, or upstream database records (uni, 2025; O’Leary et al., 2016; Blum et al., 2021; Criel et al., 2021). Conversely, the same exact sequence may carry different functional labels, host assignments, or evidence states. Database records are therefore better treated as source observations whose relationships and evidence lineages must be resolved before consolidation into reusable biological entities (Promponas et al., 2015; Kress et al., 2023; Goudey et al., 2022). Preserving native annotations and provenance, tracking source dependencies, and distinguishing experimental, curated, annotation-based, prediction-derived, and unresolved support are particularly important for machine-learning reuse, where duplicated sequences, uncertain labels, and hidden dependencies can affect dataset composition and evaluation (Goudey et al., 2022; Bell and Lord, 2017; Wilkinson et al., 2016).

Here, we present PhageLysData, an evidence-aware and AI-ready resource for phage lytic enzymes and depolymerases. The release integrates 807,366 source observations from seven primary resources into 759,105 unique exact-sequence entities organized into an evidence-supported Core, a Prediction Extension containing prediction-only candidates, and Context entities retained for provenance and reference. The Core is enriched with harmonized biological annotations, physicochemical properties, frozen UniProtKB information, independent InterProScan-derived annotations, mapped PDB and AlphaFold DB structural assets, and numerical sequence representations spanning 11 protein language models and one-hot encoding (uni, 2025; Blum et al., 2021; Burley et al., 2025; Varadi et al., 2024). By combining exact-sequence identity, evidence-aware organization, provenance-preserving integration, and reusable computational assets, PhageLysData provides a traceable foundation for protein retrieval, comparative analysis, task-specific dataset construction, and downstream machine-learning applications.

## 2 Materials and Methods

PhageLysData was constructed through a versioned workflow that separated source acquisition, exact-sequence integration, evidence-aware biological interpretation, Core enrichment, characterization, and public release (Figure **1**). Source-specific processing was completed independently before multisource integration. Resource-universe membership, canonical target classes, and evidence tiers were then assigned and frozen before downstream computational enrichment. Within the evidence-supported Core, the tiers capture increasing differences in evidence strength and resolution, from direct experimental support to curated or independently supported evidence, annotation-based support, and ambiguous or incompletely resolved cases. All subsequent physicochemical, functional, structural, and numerical-representation layers were added as enrichment only and did not alter these frozen biological assignments.

**Figure 1:**
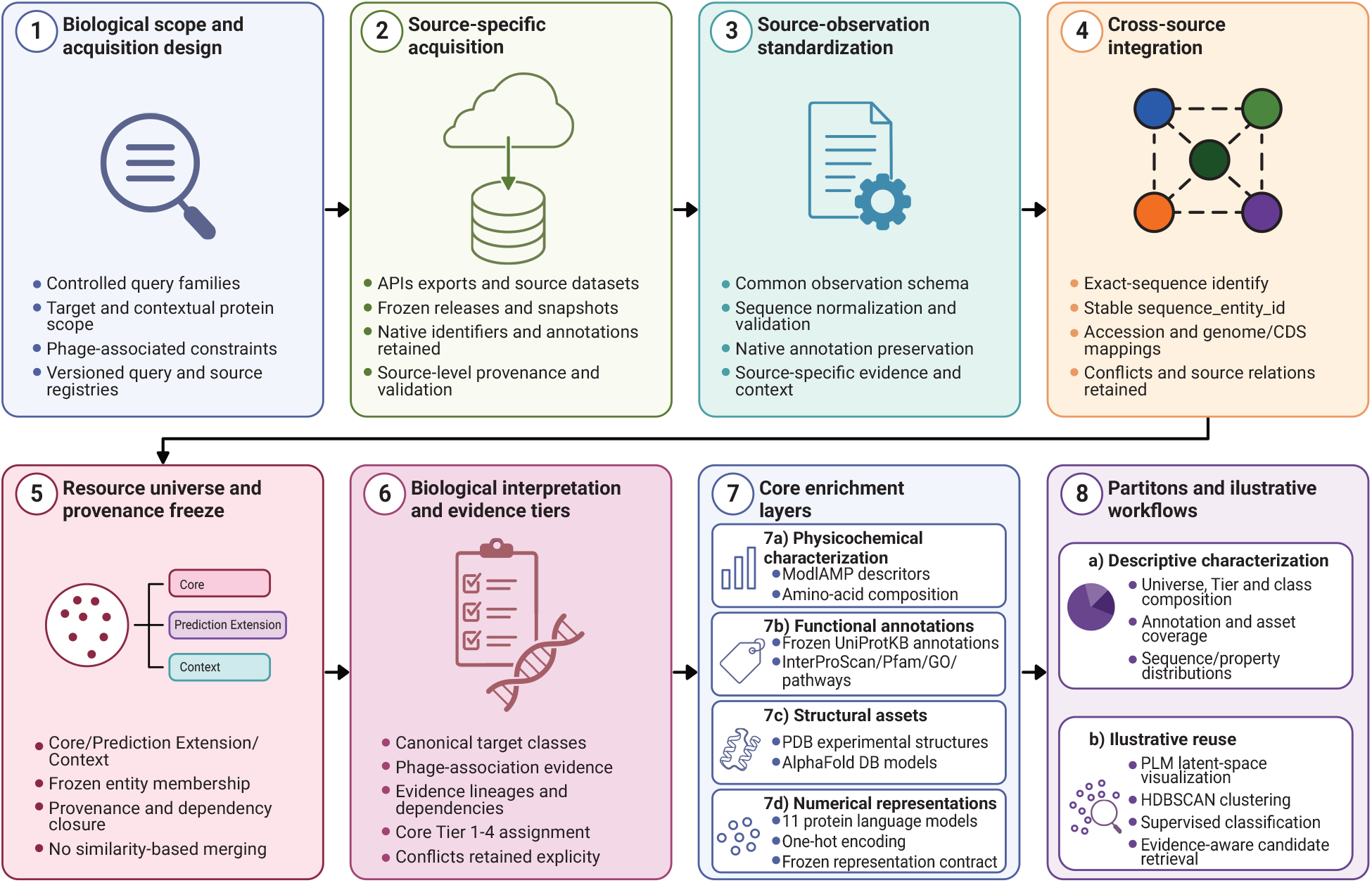
Construction and release workflow of PhageLysData. Seven primary sources were processed independently before exact-sequence integration and evidence-aware assignment of the Core, Prediction Extension, and Context universes. Core entities were subsequently enriched with physicochemical, functional, structural, and numerical representation layers without altering frozen biological assignments. Released assets remain connected through stable `sequence_entity_id` values and are accompanied by provenance, validation, and release metadata.

Stable `sequence_entity_id` values provide the common key connecting source observations, provenance, evidence, harmonized annotations, enrichment assets, and numerical representations. This design preserves the breadth of the candidate sequence space while keeping sequence identity, database occurrence, computational prediction, and biological evidence as distinct concepts.

### 2.1 Data sources and source-specific acquisition

Primary acquisition comprised seven complementary resources, including UniProtKB release 2026_02 (uni, 2025), PhaLP 2.0 (Boulay et al., 2026), INPHARED (Cook et al., 2021), DePP (Magill and Skvortsov, 2023), DposFinder (Shen et al., 2026), DepoCatalog (Otwinowska et al., 2026), and PhagesDB (Russell and Hatfull, 2017). These sources represent general protein annotation, specialized phage-enzyme collections, genome-centred resources, curated depolymerase datasets, and prediction-oriented candidate collections. Releases or snapshots, acquisition mechanisms, retrieval dates, licences, checksums, and source roles were frozen before integration. Complete source metadata and source-specific processing procedures are described in Supplementary Methods S1.

The biological scope comprised four non-exclusive target classes, including `endolysin, virion_associated_lytic_enzyme`, `depolymerase`, and `other_phage_envelope_lytic_enzyme`. Virion-associated lytic enzymes were retained as a distinct class because their definition includes physical association with the phage particle and activity during early infection, whereas `other_phage_envelope_lytic_enzyme` captures supported envelope-lytic proteins for which the available evidence does not justify assignment as an endolysin, depolymerase, or virion-associated enzyme. Broader lysis-system components, ambiguous structural proteins, and other phage proteins were retained when required to preserve provenance or biological context but were not automatically considered target entities.

UniProtKB candidates were recovered through a frozen registry of controlled queries spanning explicit protein identities, functional and catalytic annotations, Gene Ontology (Consortium, 2019) and Enzyme Commission terms (McDonald and Tipton, 2023), substrate and structural terminology, and phage-associated context. Query provenance was preserved independently from biological evidence so that synonymous or overlapping retrieval strategies did not increase confidence by themselves. The remaining resources were processed according to their native data models, with source-specific rules distinguishing curated annotations, source-reported experimental support, imported annotations, computational predictions, genomic context, and records used only as computational background. Detailed query design, retrieval procedures, and interpretation rules for each primary resource are provided in Supplementary Methods S1.

### 2.2 Exact-sequence integration and evidence architecture

Source records were converted into a common observation layer while retaining native identifiers, sequences, annotations, phage and host information, biological assertions, evidence attributes, prediction status, and source provenance. Amino-acid sequences were normalized before integration, and non-canonical residues were retained as quality attributes without triggering automatic exclusion. Detailed observation fields and normalization rules are provided in Supplementary Methods S2.

Exact amino-acid sequence identity defined the entity layer. Records corresponding to the same normalized sequence were consolidated under a stable `sequence_entity_id`, while alternative accessions, genome occurrences, phages, hosts, publications, and source-specific annotations remained connected through relation tables. Sequence similarity, homology, domain architecture, predicted function, and clustering were not used to merge entities.

After multisource integration, biological assertions were harmonized to the four target classes and linked to their provenance and evidence lineages. Core entities supported by more than one non-predictive target class were retained under the derived resource-level label `multiple_nonexclusive` and were not forced into a single canonical target class. Conflicting or discordant assertions were preserved, and repeated occurrence across databases was not interpreted automatically as independent support. Entities were then assigned to the mutually exclusive Core, Prediction Extension, or Context universes according to whether valid target support was non-predictive, exclusively prediction-derived, or absent, respectively. Complete entity-construction rules, provenance relations, universe predicates, and conflict-handling procedures are provided in Supplementary Methods S2.

### 2.3 Core enrichment and numerical representations

All frozen Core entities were retained for enrichment irrespective of whether a particular downstream annotation or asset could be obtained. Physicochemical characterization was performed with modlAMP 4.3.2 (Müller et al., 2017) and included canonical amino-acid composition, global sequence properties, and full-sequence means derived from established amino-acid scales. Descriptor incompatibility did not remove sequences from the Core; unavailable chemistry-dependent values were retained as missing with explicit status. Full descriptor definitions, parameters, and missing-value semantics are provided in Supplementary Methods S3.

A frozen UniProtKB annotation layer was materialized from release 2026_02 without performing live queries during enrichment. Accessions were connected to Core entities through exact sequence identity, retaining protein and gene annotations, taxonomy, review and annotation status, sequence features, publications, and normalized cross-references. When several UniProtKB accessions mapped to the same exact sequence, a representative accession was selected deterministically while all alternative mappings remained available. UniProt-reported external cross-references were retained as UniProt-derived information and were not treated as independently retrieved evidence.

Independent functional enrichment was performed with InterProScan 5.78-109.0 using InterPro release 109.0 (Jones et al., 2014) and Pfam release 38.2 (Mistry et al., 2021). Resulting member-database signatures, InterPro entries, Pfam assignments, domain coordinates, Gene Ontology mappings, and pathway mappings were connected to `sequence_entity_id` and stored independently from corresponding UniProtKB-reported cross-references. InterProScan annotations were treated as computational enrichment and did not alter Core membership, canonical target class, or evidence Tier. Environment validation, input preparation, execution parameters, and output materialization are described in Supplementary Methods S3.

Existing structural assets were mapped from PDB (Burley et al., 2025) and AlphaFold DB (Varadi et al., 2024). PDB relations combined frozen UniProtKB cross-references with exact sequence matching against PDB sequence records, whereas AlphaFold DB models were accepted only when their reported sequence matched the corresponding Core entity. Experimental PDB structures and AlphaFold DB models remained distinct asset classes. No additional structure prediction was performed for PhageLysData v1.0. Detailed mapping, retrieval, validation, and structural-status rules are provided in Supplementary Methods S3.

Numerical representations were generated for Core sequences containing only the 20 canonical amino acids and no more than 1,024 residues. Eleven pretrained protein language models spanning Ankh (Elnaggar et al., 2023), ESM-C (Candido et al., 2026), ESM-2 (Lin et al., 2022), Mistral-Prot (Mourad, 2024), ProtBERT (Elnaggar et al., 2021), and ProtT5 (Elnaggar et al., 2021) were executed through Sylphy under a common representation contract (Medina-Ortiz et al., 2026). Final hidden-layer representations were aggregated using mean sequence pooling and stored in FP32 precision. A position-specific one-hot representation was generated for the same eligible entity set using 20 amino-acid channels and a maximum sequence length of 1,024 residues. Model checkpoints, output dimensions, tokenization and execution metadata, and one-hot encoding details are provided in Supplementary Methods S3.

### 2.4 Resource closure, validation, and public release

Resource construction was frozen only after the Core, Prediction Extension, and Context assignments and designated Core enrichment layers had passed their corresponding validation contracts. Missing annotations, structural assets, or numerical representations were treated as availability states and did not alter an entity’s resource universe, canonical class, or evidence Tier. After construction closure, descriptive characterization summarized resource composition, evidence and class distributions, sequence properties, annotation coverage, structural availability, and representation completeness without generating new biological assignments or modifying the frozen resource.

Validation was applied throughout acquisition, integration, universe assignment, enrichment, characterization, and release packaging. Controls included source and checksum integrity, sequence and hash consistency, entity uniqueness and closure, observation-to-entity relations, evidence and provenance consistency, mutually exclusive resource-universe assignment, Tier predicates, enrichment mappings, numerical integrity, representation dimensions and row alignment, structure-file integrity, and final release inventory closure. Blocking validation failures prevented progression to downstream stages, whereas non-blocking limitations and unavailable assets were retained with explicit status and reason fields. The complete validation framework and stage-level closure criteria are described in Supplementary Methods S4.

The public PhageLysData v1.0 package was materialized exclusively from frozen construction and characterization outputs. It includes Core, Prediction Extension, and Context tables; sequences and physicochemical properties; UniProtKB and InterProScan-derived annotation layers; PDB and AlphaFold DB assets; protein-language-model and one-hot representations; characterization outputs; and machine-readable meta-data. Schemas, processing contracts, manifests, checksums, provenance records, and validation summaries accompany the release, while large immutable assets remain linked through `sequence_entity_id` rather than duplicated across tables. Release organization and asset-level metadata are detailed in Supplementary Methods S4.

### 2.5 Illustrative reuse workflows

Four release-facing workflows were implemented to demonstrate direct reuse of the distributed assets without defining a universal benchmark. Representation-space exploration used the released ESM-2 t30 150M embeddings with PCA followed by UMAP, with biological classes and evidence tiers added only as post-projection annotations. Unsupervised exploration applied HDBSCAN to the PCA-reduced representation space and used UMAP solely for visualization of the resulting computational clusters.

A supervised example evaluated Logistic Regression, linear Support Vector Machine, and Random Forest classifiers using ESM-2 t30 150M representations of four demonstration classes: depolymerase, endolysin, virion-associated lytic enzyme, and an `other` group combining `other_phage_envelope_lytic_enzyme` and `multiple_nonexclusive`. A single stratified train–test split was used without homology-aware separation or hyperparameter optimization; consequently, the analysis was interpreted as a demonstration of data and representation interoperability rather than a benchmark of remote-sequence generalization.

A final evidence-aware retrieval example combined canonical class, evidence Tier, and functional-annotation availability to select high-support depolymerases and inspect their associated numerical and structural assets. This workflow performed deterministic filtering only and did not rank candidates or predict biological desirability. Complete dimensionality-reduction, clustering, classification, and candidate-retrieval parameters are provided in Supplementary Methods S5 and the accompanying executable notebooks.

## 3 Results

### 3.1 Resource composition and evidence architecture

The final PhageLysData release integrates 807,366 source observations from seven primary resources into 759,105 unique exact-sequence entities (Figure **2**A). The difference between observations and entities reflects repeated exact amino-acid sequences occurring across accessions, genomes, source-specific identifiers, and databases while retaining their individual provenance and biological context.

**Figure 2:**
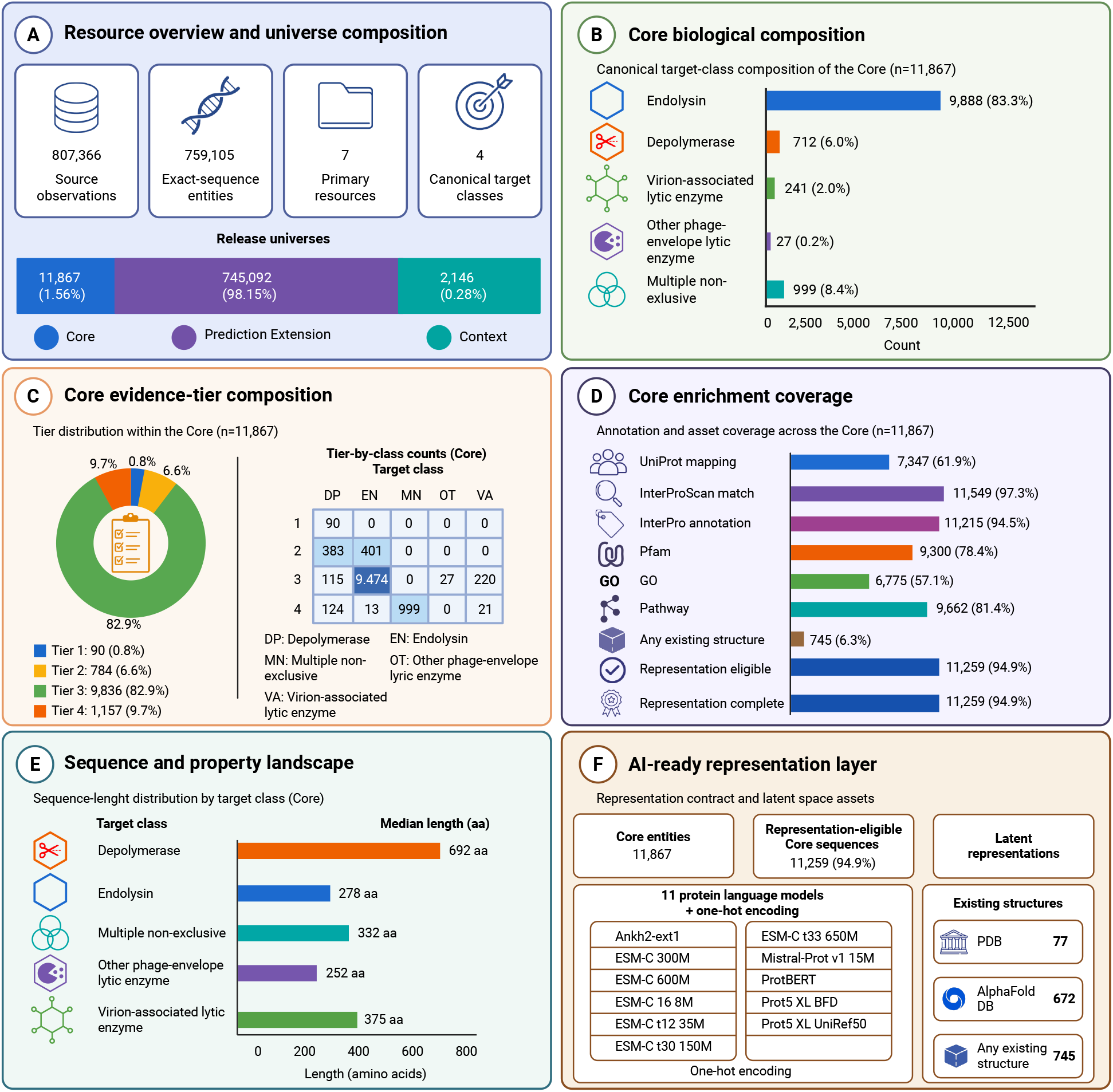
Composition and computational enrichment of PhageLysData. (A) Distribution of the 759,105 unique exact-sequence entities among the Core, Prediction Extension, and Context universes. (B) Canonical biological-class composition of the Core. (C) Core evidence-tier composition and Tier-by-class distribution. (D) Functional, structural, and numerical asset coverage. (E) Median sequence length across canonical Core classes. (F) Numerical representation and existing structural assets. Percentages in panels B–F use the frozen Core or the explicitly indicated eligible subset as denominator.

The integrated sequence space is substantially broader than the evidence-supported subset. The Prediction Extension contains 745,092 entities (98.15%) whose valid target assertions are exclusively prediction-derived, whereas the Core comprises 11,867 entities (1.56%) with at least one non-predictive target assertion. A further 2,146 entities (0.28%) are retained as Context because no valid target assertion supported assignment to the target protein space. These mutually exclusive universes preserve the breadth of the retrieved candidate space without treating prediction-derived or contextual records as equivalent to evidence-supported target entities. Prediction-derived information associated with Core entities remains available alongside non-predictive assertions, allowing concordant and discordant support to be inspected explicitly. Detailed provenance and source-richness characterization is provided in Supplementary Results S1.

Within the Core, endolysins represent the predominant canonical class, comprising 9,888 entities (83.3%), followed by 999 entities with multiple non-exclusive assignments (8.4%), 712 depolymerases (6.0%), 241 virion-associated lytic enzymes (2.0%), and 27 other phage-envelope lytic enzymes (0.2%) (Figure **2**B). Evidence support was dominated by Tier 3, with 9,836 entities (82.9%), whereas Tier 1, Tier 2, and Tier 4 contained 90 (0.8%), 784 (6.6%), and 1,157 (9.7%) entities, respectively (Figure **2**C). Tier 1 consisted exclusively of depolymerases, while most entities carrying multiple non-exclusive target assignments remained in Tier 4. Complete Tier-by-class distributions are provided in Supplementary Results S2.

### 3.2 Core enrichment and AI-ready computational assets

The frozen Core was characterized and enriched without altering its biological assignments. UniProtKB accessions were mapped by exact-sequence identity for 7,347 entities (61.9%), while independent InterProScan analysis produced at least one match for 11,549 entities (97.3%) (Figure **2**D). InterPro annotations were available for 11,215 entities (94.5%), Pfam assignments for 9,300 (78.4%), Gene Ontology mappings for 6,775 (57.1%), and pathway mappings for 9,662 (81.4%). Detailed InterProScan, Pfam, Gene Ontology, pathway, and class-specific annotation statistics are reported in Supplementary Results S3.

Existing structural coverage was more limited. PDB assets were mapped to 77 Core entities and AlphaFold DB models to 672, yielding 745 entities (6.3%) with at least one existing structural asset after accounting for overlap (Figure **2**D,F). Structural availability was retained as an independent enrichment layer and did not affect Core membership.

The Core also spans distinct sequence-length regimes among biological classes (Figure **2**E). Depolymerases were the longest class, with a median length of 692 amino acids, compared with 375 for virion-associated lytic enzymes, 332 for multiple non-exclusive entities, 278 for endolysins, and 252 for other phage-envelope lytic enzymes. Additional sequence-length and physicochemical distributions are provided in Supplementary Results S2.

Of the 11,867 Core entities, 11,259 (94.9%) satisfied the common representation contract of canonical amino acids and a maximum length of 1,024 residues. All eligible entities received the complete numerical representation layer, with no loss between eligibility and final representation materialization (Figure **2**D,F). The release therefore provides 11 protein language model embedding spaces spanning Ankh, ESM-C, ESM-2, Mistral-Prot, ProtBERT, and ProtT5, together with one-hot encoding, over a common set of stable sequence identifiers. These representations complement physicochemical, functional, and structural information and allow the same Core entities to be analysed across multiple numerical feature spaces.

### 3.3 Illustrative reuse of the released assets

Release-facing examples were used to verify that the distributed biological annotations and numerical assets could be combined directly in conventional analytical workflows. Complete analyses and visualizations are provided in Supplementary Results S4 and the accompanying notebooks.

The ESM-2 t30 150M representation contained all 11,259 representation-complete entities as 640-dimensional vectors without missing identifiers or non-finite values. PCA and UMAP were used to explore this latent space, with canonical target classes and evidence tiers overlaid only after dimensionality reduction. HDB-SCAN applied to the corresponding PCA-reduced representation identified 53 computational clusters containing 5,688 entities, while 5,571 entities (49.5%) remained classified as noise. Several clusters showed strong enrichment for individual canonical classes, although these groupings were treated as properties of the selected representation rather than newly inferred biological families.

The same representation supported an illustrative four-class classification task comprising depolymerases, endolysins, virion-associated lytic enzymes, and a combined other class. Across Logistic Regression, linear Support Vector Machine, and Random Forest models, macro-F1 ranged from 0.806 to 0.818. Because this example used a conventional stratified train–test split without homology-aware separation or hyperparameter optimization, these values demonstrate interoperability of the released class definitions and representation matrices rather than remote-sequence generalization.

Evidence-aware filtering provided a complementary reuse example that did not depend on predictive modelling. Among 712 canonical depolymerases, 473 belonged to Tier 1 or Tier 2, and progressive requirements for InterPro and Pfam support yielded 349 high-support entities. Of these, 304 had complete numerical representations and 14 had an existing mapped structure. This workflow illustrates how evidence, functional annotations, representations, and structural availability can be combined to construct task-specific subsets while retaining their underlying provenance.

## 4 Data Value and Reuse Potential

PhageLysData reduces the effort required to reconstruct provenance, evidence relationships, and computational assets when assembling datasets of phage lytic enzymes and depolymerases. Stable `sequence_entity_id` values connect native source observations with harmonized biological assignments, evidence lineages, phage and host context, functional annotations, physicochemical properties, structural assets, and numerical representations. The explicit separation of evidence-supported, prediction-derived, and contextual sequence space allows users to define task-specific inclusion criteria without discarding the provenance needed to inspect conflicting annotations or reinterpret biological support. The same entity framework also enables direct combination of complementary biological and numerical layers for candidate retrieval, comparative annotation, dataset construction, representation-space analysis, and machine-learning applications.

The resource should not be interpreted as a uniformly validated collection of phage enzymes. Prediction Extension entities remain computational candidates, some Core records retain ambiguous or incompletely resolved evidence, and missing annotations or structural assets do not imply biological absence. Likewise, the released analytical workflows demonstrate interoperability and reuse but do not constitute homology-controlled benchmarks or independent biological validation. PhageLysData is distributed as a versioned, machine-readable resource with schemas, contracts, manifests, checksums, provenance records, validation reports, and executable examples. This architecture allows future releases to incorporate revised evidence, additional sources, updated annotations, structural coverage, or new numerical representations while preserving the exact-sequence identity and provenance framework on which downstream datasets are constructed.

## Code and Data Availability Statement

The complete, versioned PhageLysData v1.0 resource is archived on Zenodo under DOI https://doi.org/10.5281/zenodo.22046027.

PhageLysData-original components, including the harmonization framework, controlled vocabulary, meta-data schema, evidence-resolution rules, documentation, and derived records, are distributed under the Creative Commons Attribution 4.0 International (CC BY 4.0) license. Third-party sequences, annotations, structural assets, and other source-derived content remain governed by the terms and licences of their original providers.

Code, workflows, executable notebooks, and environment specifications are available at https://github.com/kren-ai-lab/phagelysdata under the MIT License.

## Author Contributions Statement

AO-N, JR, and DM-O: conceptualization. R.O and DM-O: methodology. DM-O: validation. AO-N, JR, MEL and DM-O: investigation. AO-N, JR, MEL and DM-O: writing and editing. JR and MEL: supervision and funding resources. JR: project administration. All authors reviewed and approved the final version of the manuscript.

## Acknowledgments

This research was funded by the Chilean National Agency for Research and Development (ANID), through FONDECYT Exploración Grant No. 13240071 and Anillos Regulares de Tecnología Grant No. ACT240045.

## AI Statement

As non-native English speakers, the authors used ChatGPT (OpenAI, GPT-5.6 Thinking; accessed July 2026) for English editing, clarity, and consistency. The authors critically reviewed and edited all outputs, verified the final text, and take full responsibility for the manuscript.

