## Supplementary Information for "PhageLysData: an evidence-aware and AI-ready dataset of phage lytic enzymes and depolymerases"

#### Supplementary contents

**S1 Supplementary Methods** **2**

### S1 Supplementary Methods

The Supplementary Methods provide the implementation-level detail underlying the resource-construction framework described in the main text. PhageLysData was developed as a sequence of source-specific, integration, ontology, enrichment, representation, validation, characterization, and release stages. Each stage consumed frozen upstream assets and generated its own machine-readable validation, provenance, manifest, and handoff records. Unless explicitly stated otherwise, source-specific stages did not perform cross-source consolidation, and downstream enrichment stages did not modify exact-sequence identity, resource-universe membership, canonical target class, or evidence Tier.

#### S1.1 Primary sources, acquisition, and source-specific processing

##### S1.1.1 Source registry and frozen snapshots

Seven primary resources were integrated into PhageLysData: UniProtKB release 2026_02 [1], PhaLP 2.0 [2], INPHARED [3], DePP [4], DposFinder [5], DepoCatalog [6], and PhagesDB [7]. Sources were processed independently before integration so that native identifiers, source-specific evidence semantics, prediction status, and upstream dependencies could be retained without being prematurely harmonized (Table **S1**).

For each source, the acquisition layer recorded the source product, release or snapshot identity when available, native input assets, acquisition mechanism, retrieval or processing date, file size, SHA-256 checksum, licence information when available, and the source role within PhageLysData. Programmatically retrieved resources additionally retained request or batch-level checkpoints. Locally distributed packages, supplementary files, database dumps, and repository snapshots were checksum-frozen before biological interpretation.

The seven sources were intentionally treated as complementary rather than equivalent evidence providers. UniProtKB supplied a broad query-derived protein universe and rich database annotations; PhaLP supplied a specialized phage-lysin resource with source annotations and several computational layers; INPHARED and PhagesDB provided genome-centred phage context; and DePP, DposFinder, and DepoCatalog supplied focused depolymerase collections with different evidence models. These differences were preserved through source-specific evidence categories and were reconciled only after multisource integration.

**Supplementary Table S1**: **Primary PhageLysData sources and their roles in resource construction.**

| Source | Frozen acquisition unit | Primary role | Key source-specific treatment |
|---|---|---|---|
| UniProtKB | Release 2026_02; query memberships and accession hydration | Broad protein retrieval and annotation | Controlled query registry; query-level provenance retained; overlapping queries consolidated through predefined evidence-independence groups |
| PhaLP 2.0 | Frozen SQL dump and extracted native tables | Specialized phage-lysin resource | Native names, SUBLYME predictions, GO/EC, domains, SPAED, structures, literature, and contextual relations retained as distinct assertion families |
| INPHARED | Frozen genome-centred source package and annotation tables | Phage-genome and annotation context | Deterministic controlled annotation rules; GenBank- and Prokka-derived annotations treated as the same lineage when representing the same underlying annotation |
| DePP | Frozen repository, training matrix, and hydrated positive records | Depolymerase candidate source | Positive training members retained as source-curated depolymerase candidates; ML background records retained outside target evidence |
| DposFinder | Frozen curated source package | Curated depolymerase source | Source-reported experimental flag preserved without converting absence of the flag into negative evidence |
| DepoCatalog | Frozen authoritative supplementary catalog | Curated and experimentally tested depolymerases | Proteins tested in the source study distinguished from literature-derived catalog entries |
| PhagesDB | Frozen phage catalog plus available GenBank and local submission annotations | Genome-centred phage protein source | Only direct lytic/depolymerase functional annotations promoted to target candidates; generic structural or lysis-system annotations retained as context |

##### S1.1.2 UniProtKB query design and acquisition

UniProtKB acquisition was designed as a controlled retrieval experiment. The final registry contained 138 non-zero queries frozen against UniProtKB release 2026_02. Queries covered explicit target-protein names, catalytic and functional descriptions, Gene Ontology annotations [8], Enzyme Commission terms [9], substrate-related terminology, structural terminology, and phage-associated biological context.

Each query was assigned one of four retrieval depths describing its intended specificity (Table **S2**). D1 represented explicit target identity, D2 function-specific retrieval, D3 context-supported retrieval, and D4 exploratory retrieval. The final registry contained 20 D1, 71 D2, 24 D3, and 23 D4 queries. Retrieval depth was an acquisition attribute only and was never interpreted as a final evidence Tier.

**Supplementary Table S2**: **UniProtKB retrieval-depth system.** Retrieval depths describe query specificity and are distinct from the final Core evidence Tier system.

| Depth | Interpretation | Queries | Use |
|---|---|---|---|
| D1 | Explicit identity | 20 | Direct terminology identifying a target enzyme or highly specific biological identity |
| D2 | Function specific | 71 | Catalytic or functional terminology strongly associated with the target biological space |
| D3 | Context supported | 24 | Functional terminology requiring additional biological or phage-associated context |
| D4 | Exploratory | 23 | Broad structural, contextual, or discovery-oriented retrieval intended to increase recall |

The query registry was audited before sequence retrieval. Zero-hit formulations were diagnosed separately to distinguish biologically valid zero-result queries from terminology or taxonomic formulations that were unnecessarily restrictive. Rescue queries were generated deterministically when appropriate, while redundant or biologically overbroad rescue formulations were retired. The final execution plan therefore contained only queries returning at least one record within the same frozen UniProtKB release.

Each query retained its complete expression, target concept, retrieval depth, taxonomic strategy, lineage to earlier query formulations, and an `independence_group`. The latter grouped synonymous or closely overlapping retrieval formulations representing the same biological search concept. Query overlap was therefore preserved as provenance but did not automatically constitute multiple independent evidence units.

Query memberships were retrieved using UniProtKB cursor pagination with up to 500 records per request. Raw page responses, request checkpoints, query completion states, expected record counts, and query-level manifests were retained. Retrieved memberships were consolidated into a normalized query–accession relation table, while the union of unique accessions defined the hydration universe.

Each unique accession was hydrated once using 58 UniProtKB fields. Hydration included canonical amino-acid sequence, protein and gene names, reviewed status, annotation score, protein existence, taxonomy, sequence features, functional annotations, publications, virus–host information, Gene Ontology, EC and Rhea identifiers, and selected external database cross-references. Raw hydration responses were retained and normalized into accession-level and relation-level tables. Query-membership retrieval and accession hydration were locked to the same UniProtKB release; execution was aborted if release consistency could not be established.

At this stage, every query–accession relation remained traceable. Source-specific UniProt evidence units were subsequently consolidated within `independence_group` so that multiple lexical variants of the same retrieval concept did not inflate evidence multiplicity. Preliminary UniProt grades `U-D1`–`U-D4` reflected the strongest retrieval depth supporting a target assertion and remained source-specific descriptors rather than final Core evidence Tiers.

##### S1.1.3 Source-specific processing rules

The remaining six sources were interpreted according to their native data models. Their records were not forced into a common evidence vocabulary before the native biological meaning of each source field had been resolved.

**PhaLP 2.0.** The PhaLP SQL dump was first checksum-frozen and audited structurally. Table definitions, keys, relationships, row counts, and high-value fields were reconstructed directly from the SQL package before native tables were extracted into normalized intermediate assets.

PhaLP assertions were separated by origin. The native `lysin_proba` values and functional `type` assignments were interpreted as SUBLYME-derived computational predictions and assigned the evidence category `P-PRED`. Protein names were treated as source annotations (`P-NAME`) and did not constitute experimental evidence. Imported GO, EC, and related annotations were represented as database-derived annotations (`P-DBA`), while automatic annotation systems were retained separately as rule-based annotations (`P-RULE`).

Domain records were assigned `P-DOM`; SPAED-derived architecture assignments were recorded as `P-SPAED`; PDB relations as `P-STR`; bibliographic links as `P-LIT`; phage, host, cluster, and coding-sequence information as `P-CTX`; and external identifiers as `P-XREF`.

An EC assertion received `P-EXP` only when the native PhaLP evidence field explicitly reported experimental support. PubMed, DOI, or PDB linkage alone did not create experimental evidence. Assertions imported from UniProtKB were marked as shared with the UniProt lineage, whereas PhaLP-specific predictions or native information could be marked as independent from the UniProt acquisition. This distinction prevented imported annotations from being counted as independent cross-source confirmation.

**INPHARED.** INPHARED genome and annotation records were normalized before candidate selection. Candidate identification used a frozen controlled registry of annotation rules applied to functional fields including gene, product, note, function, inference, standard name, database cross-reference, EC number, and COG information.

Explicit terminology such as *endolysin*, *Lysin A*, phage lysozyme, peptidoglycan hydrolase, muralytic enzyme, muramidase, glucosaminidase, peptidoglycan amidase, peptidoglycan endopeptidase, and lytic transglycosylase supported endolysin candidate assertions according to predefined rule strength. Explicit *depolymerase*, capsule/polysaccharide lyase or hydrolase, and endosialidase terminology supported depolymerase assertions. Explicit combinations of virion-, tail-, or baseplate-associated terminology with muralytic activity supported `virion_associated_lytic_enzyme`. Lysin B or mycolylarabinogalactan esterase annotations were assigned to `other_phage_envelope_lytic_enzyme`.

Generic holin, antiholin, pinholin, spanin, lysis-system, or lysis-cassette annotations were retained as lysis-system context and did not define target enzymes. Tail-spike annotations without explicit lytic or depolymerase activity were retained as structurally ambiguous context. Candidate-rule grades `I-D1`–`I-D4` represented decreasing source-specific annotation specificity and did not correspond to final resource Tiers.

All INPHARED annotations were considered database- or computationally derived unless explicit evidence supported a stronger interpretation. No annotation was treated as experimental by default. GenBank feature annotations and auxiliary Prokka annotations were assigned to the same evidence lineage when feature identity, annotation type, and normalized value indicated that they represented the same underlying annotation.

**DePP.** The DePP repository and its training assets were frozen before candidate extraction. The authoritative training matrix was used to separate 50 positive training members from 49 machine-learning background records. Positive records were hydrated through NCBI Protein using accession.version identifiers to recover exact sequences and contextual metadata.

Membership in the positive DePP training class was interpreted as a source-curated assertion that DePP reports the sequence as an experimentally active depolymerase. This assertion was not upgraded to a manually verified primary experiment. All positive candidates used in the current source layer therefore received `D-D2`, defined as a DePP-positive sequence linked to its training membership and NCBI Protein record without independent verification of the primary experiment. `D-D1` was reserved for a directly verified primary publication and was unused in the source layer. `D-D3` was reserved for computational prediction/evaluation candidates, and `D-D4` for ML background or contextual records.

The 49 background members were excluded from the target-candidate evidence layer and were not interpreted as biological negatives. NCBI Protein supplied identity, phage context, host qualifiers, coding information, and bibliographic links but did not constitute an independent depolymerase assertion.

**DposFinder.** The DposFinder source layer contained 384 curated depolymerase records with resolved sequence information. The source field `experimental=yes` was interpreted strictly as source-reported experimental support and was not promoted to a manually verified primary experiment. These records received `DF-D2`. Records curated as depolymerases without an experimental flag received `DF-D3`; `experimental=no` did not represent functional negative evidence.

The released source layer contained 103 `DF-D2` and 281 `DF-D3` observations. DOI identifiers were retained as publication provenance and used to construct bibliographic evidence lineages. Target receptor, serotype, and domain fields were preserved, including explicit unknown values. `target_receptor=unknown` and missing experimental flags were treated as missing source information, not as biological negatives.

**DepoCatalog.** DepoCatalog records were extracted from the authoritative supplementary catalog. Exact protein sequences, phage information, target information, source-study testing status, structural annotations, and bibliographic provenance were normalized without website scraping or reinterpretation of external structural models.

Two source-specific evidence grades were used. `DC-D1` identified proteins for which the exact sequence and target were reported and the protein was tested in the current DepoCatalog study. `DC-D2` identified exact sequence–target records derived from prior literature but not tested in the current study. Source inconsistencies were retained explicitly, including documented sequence-level conflicts, instead of being silently corrected during normalization.

**PhagesDB.** The frozen PhagesDB package comprised the catalog of sequenced phages and all available annotation assets. Public GenBank records were downloaded and cached for phages with accessions, while finalized local NCBI submission files were parsed when public accessions were unavailable. Phages lacking either annotation source remained represented at the phage level with explicit annotation-unavailable status.

Protein candidates were selected using a deliberately high-precision annotation policy. Direct product annotations reporting a lytic enzyme, peptidoglycan-active enzyme, lysin, lysozyme, muramidase, endolysin, depolymerase, or defined polysaccharide-degrading activity were sufficient for target promotion. Generic *holin*, *spanin*, *lysis protein*, *tail fiber*, *tail spike*, or *baseplate* annotations were insufficient in the absence of explicit lytic or depolymerase function and were retained separately for contextual review.

PhagesDB and associated GenBank/submission annotations were treated as source-reported functional annotations rather than manually verified primary experiments. Selected candidates therefore received the source-specific grade `PHDB-D3`.

**Supplementary Table S3**: **Summary of source-specific evidence interpretation.** Source-specific grades are retained for provenance and are not equivalent to final PhageLysData evidence Tiers.

| Source | Principal source-specific grades | Interpretation | Important restriction |
|---|---|---|---|
| UniProtKB | `U-D1`–`U-D4` | Strongest query-retrieval depth supporting the target assertion | Retrieval specificity is not a final evidence Tier |
| PhaLP | `P-PRED`, `P-NAME`, `P-DBA`, `P-RULE`, `P-DOM`, `P-SPAED`, `P-EXP`, `P-LIT`, `P-CTX`, `P-STR`, `P-XREF` | Assertion origin retained explicitly | Imported UniProt information is not independent PhaLP evidence |
| INPHARED | `I-D1`–`I-D4` | Controlled annotation specificity and context | Annotation is non-experimental by default |
| DePP | `D-D2` used; `D-D1`, `D-D3`, `D-D4` reserved | Positive training membership is source-curated support | Background membership is not functional negative evidence |
| DposFinder | `DF-D2`, `DF-D3` | Source-reported experimental flag versus curated depolymerase record without such flag | Experimental flag is not independent primary-study verification |
| DepoCatalog | `DC-D1`, `DC-D2` | Tested in current study versus prior-literature catalog entry | Source-reported inconsistencies are retained |
| PhagesDB | `PHDB-D3` | Direct source product annotation of lytic/depolymerase activity | Generic structural or lysis-system terminology alone is insufficient |

#### S1.2 Exact-sequence integration, provenance, and evidence model

##### S1.2.1 Source-observation schema

The seven closed source layers were converted into a common source-observation schema during multisource integration. The observation layer was designed to preserve source-native biological meaning while exposing a minimal shared set of fields required for exact-sequence consolidation and evidence aggregation.

Each observation received a stable global observation identifier generated from the source family, frozen source snapshot, and native observation identifier. Identifiers use the `PLDOBS` prefix and remain distinct from exact-sequence entity identifiers.

The common schema retained source identity, source snapshot, native record and protein identifiers, sequence, sequence hash, sequence length, accession information, annotation text, phage and host context, candidate role, source-reported and harmonized target classes, source-specific grade, deployment, substrate target, catalytic activity, experimental-evidence flags, prediction status, and the originating source asset.

Amino-acid sequences were normalized by removing whitespace and converting residues to uppercase. Empty sequences were retained as source observations but could not create exact-sequence entities. Non-canonical or extended residue symbols were recorded through explicit quality fields and did not trigger automatic exclusion from the observation layer.

**Supplementary Table S4**: **Principal fields in the integrated source-observation schema.**

| Field group | Representative fields |
|---|---|
| Observation identity | `global_source_observation_id`, `source_family`, `source_product`, `source_snapshot_id`, `native_observation_id`, `native_record_id`, `native_protein_id` |
| Sequence identity | `sequence`, `sequence_sha256`, `sequence_length`, `sequence_valid_for_entity`, `noncanonical_residues` |
| Protein accessions | `protein_accession`, `protein_accession_normalized`, `protein_accession_base`, `accession_namespace` |
| Biological context | `annotation`, `phage_name`, `phage_accession`, `host_name`, `deployment`, `substrate_target`, `catalytic_activity` |
| Target interpretation | `candidate_role`, `source_target_class`, `harmonized_target_class`, `source_specific_grade` |
| Evidence status | `primary_experiment_verified`, `direct_experimental_evidence`, `is_prediction`, `prediction_derived_observation`, `observation_evidence_origin` |
| Provenance | `source_observation_layer`, `source_asset` |

##### S1.2.2 Exact-sequence entity model

Molecular identity was defined exclusively by complete normalized amino-acid sequence. A SHA-256 digest was calculated from each non-empty normalized sequence, and all observations with the same digest were connected to one exact-sequence entity. The stable entity identifier was constructed as the prefix `PLDSEQ_` followed by the complete sequence SHA-256 digest.

The canonical exact-sequence table contained only `sequence_entity_id`, `sequence_sha256`, amino-acid sequence, and sequence length. Biological class, evidence Tier, resource universe, prediction-only status, and exclusion status were deliberately absent from this identity table.

No pairwise sequence similarity, identity threshold, homology search, protein-domain architecture, functional label, predicted class, or clustering result contributed to entity merging. Consequently, proteins differing by even one residue remained distinct entities, while exact sequences occurring under multiple accessions, phages, genomes, source records, publications, or annotations were represented once at the entity layer and many times through relation tables.

An observation-to-entity table retained every exact mapping. A separate source-support summary recorded the number of contributing observations and source families, presence of prediction-derived observations, and source-level experimental flags without using these fields to redefine molecular identity.

##### S1.2.3 Provenance and evidence lineages

PhageLysData distinguishes source occurrence, evidence unit, and evidence lineage. A source occurrence indicates that a record was observed in a particular source. An evidence unit represents a specific assertion or set of assertions that can be treated as one support unit within that source. An evidence lineage describes upstream dependence among evidence units.

This distinction was required because two apparent source records may derive from the same upstream annotation, prediction system, or publication. Examples include multiple UniProtKB queries retrieving the same accession, PhaLP annotations imported from UniProtKB, INPHARED GenBank and Prokka representations of the same annotation lineage, multiple source entries linked to the same publication, and repeated prediction outputs generated by one computational system.

Within UniProtKB, synonymous queries belonging to the same `independence_group` were consolidated into one independent query-evidence unit. Within PhaLP, imported UniProt annotations were explicitly tagged as shared with the UniProt lineage, while PhaLP-specific SUBLYME and SPAED outputs retained their own computational lineages. INPHARED GenBank and Prokka annotations were linked when they represented the same underlying annotation. Bibliographic relations were retained as shared publication lineages when multiple observations pointed to the same DOI or PubMed identifier.

Cross-source integration therefore did not use raw source count as an evidence score. Multiple observations remained informative for provenance and context, but final evidence assignment operated on explicit assertion properties, source families, evidence units, and lineage-aware support.

###### S1.2.4 Resource-universe assignment

Resource-universe assignment was performed only after all source-level biological assertions had been integrated.

A biological assertion was considered a valid target assertion when its harmonized target-class candidate belonged to one of the four target classes: `endolysin`, `virion_associated_lytic_enzyme`, `depolymerase`, or `other_phage_envelope_lytic_enzyme`.

Prediction-derived status was inherited from individual observations and assertions. It was not inferred from the entity's source composition.

The **Core** contains entities with at least one valid non-predictive target assertion. Prediction-derived assertions can also be present for a Core entity, but they cannot override non-predictive support.

The **Prediction Extension** contains entities with at least one valid target assertion but no valid non-predictive target assertion. By definition, all valid target support for these entities is prediction-derived.

The **Context** universe contains entities for which no valid target assertion remained after integration. These sequences are retained because they preserve source, provenance, lysis-system, genome, structural, or other contextual information.

Universe assignment is therefore mutually exclusive. In formal terms, for entity $e$ with valid target-assertion set $A_e$:

$$e \in \text{Core} \quad \Longleftrightarrow \quad \exists a \in A_e \text{ such that } a \text{ is non-predictive},$$

$$e \in \text{Prediction Extension} \quad \Longleftrightarrow \quad A_e \neq \varnothing \ \wedge \ \forall a \in A_e,\ a \text{ is prediction-derived},$$

and

$$e \in \text{Context} \quad \Longleftrightarrow \quad A_e = \varnothing.$$

Prediction-derived classes were retained for Core entities and compared with the non-predictive class set. Prediction-class discordance was flagged when at least one prediction-derived class fell outside the supported non-predictive class set. Such discordance did not modify the Core assignment or replace the non-predictive canonical class.

###### S1.2.5 Canonical target classes and evidence tiers

For Core entities, canonical target class was derived only from non-predictive assertions. An entity supported by exactly one non-predictive target class received that class directly. When more than one non-predictive class remained supported, the entity was assigned the derived resource-level label `multiple_nonexclusive`. This label is not a fifth biological target class and indicates that the evidence supports more than one of the four target classes without forcing an artificial single-class resolution.

Prediction Extension entities were classified analogously from prediction-derived assertions. Entities with one predicted target class received that class, whereas entities with several prediction-derived classes received `multiple_prediction_candidates`. Context entities received no target-class assignment.

Phage association was summarized independently from target class using four levels. Level 0 represented unresolved phage association. Level 1 represented probable source context, including a target assertion without stronger phage linkage. Level 2 represented support from a phage-specialized or genome-centred source. Level 3 required an explicit entity–phage relation. Levels 2 and 3 satisfied the minimum phage-association criterion for Tier 1–3 assignment.

Evidence Tiers were assigned only within the Core. Every Core entity initially entered Tier 4 and was promoted deterministically when stronger criteria were satisfied. Prediction Extension and Context entities remained outside the Tier 1–4 system.

**Supplementary Table S5**: **Deterministic Core evidence-Tier predicates.** Criteria are evaluated from integrated non-predictive target assertions. Prediction Extension and Context entities are outside this Tier system.

| Tier | Interpretation | Deterministic requirements |
|---|---|---|
| Tier 1 | Experimentally supported | Core membership; exactly one non-predictive target class; phage-association level $\geq$ 2; no blocking conflict; `primary_experiment_verified`=true; and `direct_experimental_evidence`=true on valid non-predictive target support |
| Tier 2 | Curated high-confidence | Core membership; exactly one non-predictive target class; phage-association level $\geq$ 2; no blocking conflict; and at least one of the following: strong specialized curated support, support from at least two distinct non-general non-predictive source families, or source-reported experimental support accompanied by publication provenance |
| Tier 3 | Annotation-supported | Core membership; exactly one non-predictive target class; phage-association level $\geq$ 2; no blocking conflict; and explicit non-predictive annotation support |
| Tier 4 | Supported but ambiguous or incomplete | Core membership with non-predictive target support that does not satisfy Tier 1–3, including multiple supported target classes, insufficient phage association, weak/incomplete non-predictive support, or a blocking conflict |

Strong specialized curated support was recognized for predefined source-specific combinations from DePP, DposFinder, and DepoCatalog. Independent multisource support required at least two distinct non-general source families contributing non-predictive target assertions. UniProtKB alone could therefore not satisfy this criterion through repeated query retrieval.

Machine-readable reason codes were attached to universe, class, and Tier assignments. These codes recorded, among other conditions, non-predictive Core support, prediction-only support, single or multiple classes, concurrent prediction support, prediction-class discordance, phage-association strength, Tier 1 experimental criteria, Tier 2 curated or multisource support, Tier 3 annotation support, and Tier 4 ambiguity or conflict. Core entities requiring review because of multiple non-predictive classes, blocking conflicts, or weak phage association received an explicit review status rather than being removed.

#### S1.3 Core enrichment and numerical representations

##### S1.3.1 Physicochemical characterization

Physicochemical characterization was performed only after Core membership, canonical target class, and evidence Tier had been frozen. The analysis used modlAMP 4.3.2 [10].

Before descriptor generation, sequence length, sequence SHA-256, and the correspondence between `sequence_entity_id` and the sequence hash were revalidated. Core entities were processed in deterministic batches of 2,000 sequences.

Absolute counts and relative frequencies were calculated for each of the 20 canonical amino acids. Additional sequence-quality fields included the number of observed canonical amino-acid types, the number and fraction of non-canonical residues, and the identities of non-canonical residue types.

Chemistry-dependent descriptors were generated using `modlamp.descriptors.GlobalDescriptor.calculate_all()`. The installed modlAMP default pH of 7.4 was used and C termini were treated as non-amidated, consistent with the use of full-length proteins. Full-sequence global means were additionally calculated using the Eisenberg, GRAVY, bulkiness, and flexibility scales through `PeptideDescriptor.calculate_global()` with mean aggregation. Molecular formula was computed separately with `GlobalDescriptor.formula()`.

Chemistry-dependent descriptors were calculated only for sequences containing the 20 natural amino acids recognized by modlAMP. Sequences containing unsupported residues remained Core members; amino-acid composition and quality fields were retained, while incompatible physicochemical values and molecular formula were marked unavailable. No dataset-level scaling, normalization, imputation, or feature transformation was applied during resource construction.

Every released feature was represented in a descriptor registry containing feature name, family, implementation class, method, modlAMP version, effective parameters, eligibility rule, missing-value semantics, and inclusion status.

##### S1.3.2 Frozen UniProtKB enrichment

The Core UniProtKB enrichment layer was materialized exclusively from the upstream frozen UniProtKB 2026_02 acquisition. No live UniProtKB API queries were performed during enrichment.

Core entities were linked to frozen UniProtKB accessions through exact sequence SHA-256 identity. Accession mappings were accepted only when the sequence hash from the hydration layer agreed with the Stage-27 Core sequence hash. All accessions mapping to a Core entity were retained.

Accession-level enrichment included protein and gene names, organism and taxonomic information, reviewed status, annotation score, protein-existence information, selected sequence features, functional annotations, publication information, and normalized external database relations.

When several UniProtKB accessions mapped to one exact sequence, a representative accession was selected deterministically using the following precedence:

$$\text{reviewed} > \text{annotation score} > \text{nonfragment} > \text{accession lexical order.}$$

This representative accession was provided as a convenience field only. Alternative accessions remained available through entity–accession relation tables.

Normalized UniProtKB relations included EC, GO, Rhea, InterPro, Pfam, PDB, PubMed, RefSeq, EMBL, proteome, keyword, and virus–host identifiers. These relations were explicitly marked as UniProtKB-reported annotations or cross-references. A UniProtKB-reported InterPro, Pfam, or PDB identifier was therefore not treated as independent evidence from InterPro, Pfam, or PDB.

Absence of a frozen UniProtKB mapping indicates only that no accession from the frozen query-derived UniProt acquisition mapped to that exact sequence. It does not imply that the sequence is absent from the complete or current UniProtKB database.

##### S1.3.3 Independent InterProScan enrichment

Independent domain and functional annotation was generated using InterProScan 5.78-109.0 [11], InterPro release 109.0, and Pfam release 38.2 [12]. The production environment used Java 11.

The complete frozen Core was validated before InterProScan preparation. Input checks verified sequence length, SHA-256 identity, correspondence between `sequence_entity_id` and sequence hash, non-empty sequences, and FASTA compatibility. InterProScan protein FASTA accepted alphabetic residue symbols, while explicit forbidden symbols such as `-`, `.`, `*`, and `_` were treated as incompatible. All 11,867 Core sequences passed the production input contract.

Inputs were divided deterministically into residue-balanced FASTA shards. The number of shards was defined as the maximum of the minimum number required by a 5,000-sequence hard limit and a target residue load of 1,500,000 residues per shard. Sequences were ordered by decreasing sequence length, with `sequence_entity_id` as a stable tie-breaker, and assigned using a longest-processing-time balancing strategy to the currently lightest available shard. The final production input comprised four shards for 4,554,089 Core residues.

The runtime was preflighted before execution. Checks included the InterProScan version, InterPro release, Pfam release, Java version, available applications, installation configuration, and reachability of the configured precalculated-match lookup service.

InterProScan was executed in protein mode with TSV output, Gene Ontology mappings, pathway mappings, residue-level annotation disabled, TSV-version sidecars enabled, and eight CPUs per shard. Unless explicitly restricted, all analyses available in the frozen installation were executed. The canonical command semantics corresponded to protein mode, TSV output, `-goterms`, `-pa`, `-dra`, and `-vtsv`. Precalculated-match lookup remained enabled in the production configuration.

Each Stage-30A sequence registry additionally stored the sequence MD5 because InterProScan TSV reports MD5 values. During materialization, every raw match was validated against the expected entity identifier, MD5, sequence length, shard membership, match coordinates, signature accession, and analysis identifier.

Canonical release assets included raw member-database matches, InterPro assignments, Pfam matches and architectures, Gene Ontology mappings, pathway mappings, entity-level annotation summaries, and coverage tables. Independently observed InterPro/Pfam results were stored separately from corresponding UniProtKB-reported cross-references, allowing concordance or discordance between the two provenance layers to be examined explicitly.

##### S1.3.4 Existing structural assets

Structural enrichment was restricted to existing PDB and AlphaFold DB assets [13, 14]. No additional structure prediction was performed for PhageLysData v1.0.

PDB mappings were assembled from two routes. First, frozen UniProtKB-reported PDB cross-references associated with exact-sequence-mapped Core accessions were retained. Second, the RCSB PDB SEQRES sequence archive was scanned for sequences matching a Core sequence exactly by SHA-256. PDB IDs and chain/entity identifiers recovered from exact SEQRES matching were retained together with their mapping source and method.

Mapped PDB coordinate files were downloaded in mmCIF format and validated for non-empty, structurally valid content before registration. File size and SHA-256 were recorded.

AlphaFold DB was queried only through UniProtKB accessions previously linked to Core entities by exact sequence identity. Returned models were accepted only when the amino-acid sequence reported by AlphaFold DB matched the Core sequence exactly. Coordinate URLs and model metadata were retained, and downloaded mmCIF or PDB files were validated before inclusion.

The final structural status followed the precedence

$$\text{PDB} > \text{AlphaFoldDB} > \text{no\_existing\_structure}.$$

If a Core entity had one or more PDB structures, its preferred status was `experimental_structure`. An entity without PDB coverage but with a successfully retrieved AlphaFold DB model received `alphafold_model`. When neither coordinate asset was successfully retrieved, the entity received `no_existing_structure`. All relations were retained when both PDB and AlphaFold DB were available.

The `no_existing_structure` state is specific to the frozen retrieval workflow and does not assert universal absence of a structure or model from external databases. AlphaFold DB query or retrieval failures were retained as provenance/audit states and were not interpreted as evidence that no model exists.

##### S1.3.5 Protein language model representations

A common eligibility contract was frozen before numerical representation generation. A Core sequence was eligible when its length was no greater than 1,024 amino acids and all residues belonged to the canonical alphabet

`ACDEFGHIKLMNPQRSTVWY`.

Ineligible sequences remained unchanged Core members and retained explicit filter reasons, including `sequence_length_gt_1024` and `contains_noncanonical_amino_acid`.

Eleven pretrained protein language models were executed through Sylphy [15]. Model discovery was not dynamic; the complete model registry was frozen before execution. All PLM runs used CUDA, FP32 precision, a maximum sequence length of 1,024 residues, `last` as the selected layer, mean layer aggregation, and mean sequence pooling. Model outputs were validated for row count, entity order, sequence identity when present in the raw output, numerical finiteness, dimensional consistency, and FP32 preservation after serialization.

Each model was executed in an independent Sylphy subprocess. Requested batch sizes were conservative initial values selected for full-length sequences up to 1,024 residues; they did not alter the representation definition.

**Supplementary Table S6**: **Frozen PLM representation registry.** All models used the final hidden layer, mean sequence pooling, FP32 output, and the same representation-eligible Core population. Requested batch sizes correspond to the production Sylphy execution configuration.

| Family | Frozen model identifier | Requested batch | Representation contract |
|---|---|---|---|
| Ankh | `elnaggarlab/ankh2-ext1` | 1 | last layer; mean pooling; FP32 |
| ESM-C | `esmc_300m` | 2 | last layer; mean pooling; FP32 |
| ESM-C | `esmc_600m` | 1 | last layer; mean pooling; FP32 |
| ESM-2 | `facebook/esm2_t6_8m_ur50d` | 16 | last layer; mean pooling; FP32 |
| ESM-2 | `facebook/esm2_t12_35m_ur50d` | 16 | last layer; mean pooling; FP32 |
| ESM-2 | `facebook/esm2_t30_150m_ur50d` | 4 | last layer; mean pooling; FP32 |
| ESM-2 | `facebook/esm2_t33_650m_ur50d` | 1 | last layer; mean pooling; FP32 |
| Mistral-Prot | `raphaelmourad/mistral-prot-v1-15m` | 16 | last layer; mean pooling; FP32 |
| ProtBERT | `rostlab/prot_bert` | 2 | last layer; mean pooling; FP32 |
| ProtT5 | `rostlab/prot_t5_xl_bfd` | 1 | last layer; mean pooling; FP32 |
| ProtT5 | `rostlab/prot_t5_xl_uniref50` | 1 | last layer; mean pooling; FP32 |

Each representation was released as a typed Parquet matrix with `sequence_entity_id` as the authoritative join key and accompanied by model-specific metadata containing the model identifier, model family, extraction configuration, output dimensionality, requested batch size, precision, pooling policy, eligible-entity count, and validation status.

Representation generation was resumable at model level. A model was considered complete only when its metadata and representation matrix passed the frozen validation contract. The final representation-closure stage required all 11 PLM assets to be valid for every representation-eligible entity.

##### S1.3.6 One-hot encoding

A classical one-hot representation was generated for the same representation-eligible sequence set used by the PLMs. Eligibility was consumed directly from the PLM-stage registry and was not recalculated.

The Sylphy `one_hot` encoder used the canonical 20-amino-acid alphabet, maximum sequence length of 1,024 residues, no extended-residue channel, and no unknown-residue channel. Residues were represented positionally and sequences shorter than 1,024 residues were zero-padded after sequence termination.

The flattened representation followed position-major, residue-channel-minor ordering and contained

$$1,024 \times 20 = 20,480$$

features per sequence. Values were stored as `uint8`. The release includes the one-hot Parquet matrix, metadata, and a feature map identifying the position and residue corresponding to each flattened column.

The final numerical-representation closure required every eligible entity to possess all 12 representation assets: 11 PLM embeddings plus one-hot encoding. Filtered entities were recorded as `filtered_not_applicable` and were not treated as representation failures.

#### S1.4 Resource closure, validation, and release organization

##### S1.4.1 Construction freeze and stage closure

Construction was organized so that biological ontology and computational enrichment could not silently modify each other.

Stage 26 integrated the seven closed source layers, produced the global source-observation layer, built exact-sequence entities, and materialized provenance and evidence relations. It was not authorized to assign final resource universes or evidence Tiers.

Stage 27 was the sole authoritative stage for Core, Prediction Extension, and Context assignment, canonical target class, prediction-only status, phage-association summary, and Core evidence Tier.

Stages 28–32 enriched the frozen Core with physicochemical properties, frozen UniProtKB annotations, independent InterProScan annotations, existing structural assets, and numerical representations. These stages were explicitly prohibited from changing Stage-27 membership, class, Tier, or sequence identity.

Stage 33 performed final resource-construction closure. It generated a one-row-per-Core master table, Core FASTA, asset-availability registry, lightweight Prediction Extension and Context views, an all-entity resource index, coverage summaries, stage/asset registries, validation reports, and the final construction contract. Large PLM matrices and structural coordinate files remained immutable external assets referenced through their registries rather than being duplicated into the Core master table.

Stage 34 performed descriptive characterization only after construction closure. It summarized resource-universe composition, canonical classes, evidence Tiers, Tier-by-class composition, source/provenance richness, sequence and physicochemical distributions, annotation coverage, structural availability, and representation completeness. Stage 34 generated no new annotation, clustering, classification, predictive model, resource-universe assignment, or biological inference and did not mutate the frozen resource.

This ordering established the final construction logic:

$$
\begin{aligned}
&\text{source acquisition} \rightarrow \text{source closure} \rightarrow \text{exact sequence integration}\\
&\rightarrow \text{evidence/resource freeze} \rightarrow \text{Core enrichment}\\
&\rightarrow \text{construction closure} \rightarrow \text{descriptive characterization}\\
&\rightarrow \text{public release.}
\end{aligned}
$$

##### S1.4.2 Validation framework

Validation was implemented as a blocking component of the workflow and not solely as a final release check. Each major stage generated machine-readable validation tables or JSON records together with an explicit completion status.

Checks covered source-file integrity, manifest continuity, sequence identity, accession consistency, evidence and provenance relationships, resource ontology, enrichment integrity, representation alignment, structural file integrity, and release closure. When a check defined as blocking failed, the stage was not considered completed and its handoff was not accepted by the next stage.

Non-blocking conditions represented legitimate incompleteness or unresolved information and remained visible in explicit status fields. Examples include missing UniProtKB mappings, absence of InterPro assignments despite successful processing, unavailable existing structures, non-canonical residues preventing a subset of physicochemical descriptors, representation ineligibility, source conflicts, and unresolved contextual mappings.

A versioned contract accompanied each major post-integration stage. Contracts recorded the authoritative input stage, output scope, invariants that downstream stages were required to preserve, and the conditions under which an output was considered ready for handoff. Construction therefore depended on explicit stage closure rather than the existence of output files alone.

##### S1.4.3 Public release organization

The public PhageLysData v1.0 release was materialized from validated Stage-33 and Stage-34 outputs together with the frozen upstream enrichment assets. Release packaging was copy-only and did not mutate the construction workspace.

**Supplementary Table S7**: **Principal validation domains across PhageLysData construction.** Individual checks and observed/expected values are distributed in the corresponding stage-level validation records.

| Stage | Validation domain | Representative blocking checks |
|---|---|---|
| Source stages | Source closure | Frozen-file checksum, source-package continuity, row/identifier integrity, absence of unintended cross-source integration or automatic exclusion |
| 26 | Multisource integration | Unique `PLDOBS` identifiers, valid observation-to-entity mapping, exact sequence/hash consistency, no orphan source classifications, provenance relation closure |
| 27 | Resource ontology | Mutually exclusive Core/Prediction Extension/Context assignment, Core Tier 1–4 validity, Prediction Extension outside Core Tiers, canonical-class predicates, reason-code consistency |
| 28 | Physicochemical layer | Core identity closure, descriptor row count, finite values for compatible sequences, sequence-length agreement, descriptor-registry uniqueness |
| 29 | Frozen UniProtKB | Frozen accession/hash consistency, exact mapping to Core, representative accession availability, no live API calls, Stage-27 immutability |
| 30 | InterProScan | Core input integrity, complete shard execution, raw TSV schema, MD5/length/entity/shard consistency, valid coordinates and signatures, complete entity summary |
| 31 | Structural layer | Coordinate-file integrity, checksum agreement, relation-to-Core consistency, exact AlphaFold sequence match, final structural-status partition, Stage-27 immutability |
| 32 | Representations | Eligibility contract, exact row order, representation dimensions, FP32 PLM dtype, one-hot feature count and uint8 dtype, complete 12/12 representation coverage for eligible entities |
| 33–35 | Construction/release closure | Upstream contract compatibility, frozen resource counts, asset inventory, portable paths, absence of developer-specific absolute-path leakage, final manifest and checksum closure |

The public release was intentionally separated from raw source snapshots and the complete internal processing history. The release-facing directory contains the curated biological and computational assets required for reuse, while an organized provenance workspace can separately retain source packages and intermediate stage outputs.

The principal public structure is:

```
PhageLysData_v1.0/
|-- core/
|   |-- phagelysdata_core.parquet
|   |-- phagelysdata_core.tsv.gz
|   |-- phagelysdata_core.fasta.gz
|   |-- physicochemical/
|   `-- annotations/
|       |-- uniprot/
|       `-- interproscan/
|-- prediction_extension/
|-- context/
|-- representations/
|   |-- representation_registry.parquet
|   |-- representation_asset_catalog.tsv
|   |-- plm/
|   `-- one_hot/
|-- structures/
|   |-- coordinates/
|   |   |-- pdb/
|   |   `-- alphafold/
|   `-- ...
|-- characterization/
|-- metadata/
|   |-- contracts/
```

```
|   `-- release_asset_provenance.tsv
|-- examples/
|-- MANIFEST.tsv
`-- CHECKSUMS.sha256
```

The Core directory contains the primary entity-level table, Core FASTA, physicochemical data, and annotation relations. Prediction Extension and Context are distributed separately to prevent their use as implicit equivalents of Core entities. Representation matrices are stored independently because of their size and are joined through `sequence_entity_id`. Existing PDB and AlphaFold DB coordinates and their mapping registries are stored under the structural layer.

Release-facing metadata include the resource entity index, schemas, contracts, asset catalogs, provenance records, release metadata, and validation summaries. `release_asset_provenance.tsv` maps every public asset to its frozen construction-stage source.

A complete `MANIFEST.tsv` records relative path, release category, file size, and SHA-256 for each distributed file. `CHECKSUMS.sha256` provides a conventional checksum list for independent integrity verification.

Release portability was validated explicitly. Developer-specific absolute filesystem paths and execution commands were removed or replaced with release-relative paths in user-facing metadata. Release notebooks were required to consume files exclusively from the public release directory and were prohibited from reading internal `results/`, `raw_data/`, `processed_data/`, or developer-specific locations.

##### S1.5 Illustrative reuse workflows

The release-facing notebooks were designed as executable demonstrations of resource interoperability and not as optimized biological analyses or universal benchmarks. All examples consumed public-release assets only and did not alter the resource.

###### S1.5.1 Latent-space visualization

Latent-space visualization used the released ESM-2 t30 150M representation. The representation matrix was validated against the representation-complete Core identifier set before dimensionality reduction. Only embedding feature columns were used as numerical inputs; metadata fields were excluded from PCA and UMAP.

The embedding matrix was reduced by principal-component analysis using

$$n_{\mathrm{PCA}} = \min(50,\ n-1,\ d),$$

where $n$ is the number of sequences and $d$ the embedding dimensionality. PCA used the scikit-learn implementation with `random_state=42`.

The PCA representation was projected to two dimensions using UMAP with 30 neighbours, `min_dist=0.10`, Euclidean distance, and random seed 42. Canonical target class, evidence Tier, sequence length, annotation status, and structural availability were joined only after numerical projection and were used as visualization overlays.

The UMAP coordinates were interpreted as a descriptive visualization of one pretrained-model representation. Spatial proximity was not interpreted as evidence of homology, identical biochemical function, or a newly discovered protein family.

###### S1.5.2 Unsupervised clustering

The clustering example used the same released ESM-2 t30 150M representation and the same representation-to-Core identity checks. Clustering was performed on PCA-reduced PLM features rather than on UMAP coordinates.

At most 50 principal components were retained using the same rule defined for latent-space visualization. HDBSCAN was then applied with:

- `min_cluster_size = 50`;
- `min_samples = 15`;

- Euclidean distance;
- excess-of-mass (`eom`) cluster selection.

HDBSCAN noise assignments were retained as a valid clustering outcome. UMAP with 30 neighbours, `min_dist=0.10`, Euclidean distance, and seed 42 was generated independently from the PCA representation only for visualization of the HDBSCAN assignments.

Cluster characterization included cluster size, fraction of noise observations, canonical target-class composition, class purity, normalized class entropy, evidence-Tier composition, sequence-length summaries, and coverage of UniProtKB, InterProScan, InterPro, Pfam, GO, pathway, and existing structural assets. These summaries were descriptive and did not cause cluster-derived biological labels to be written back into PhageLysData.

##### S1.5.3 Supervised classification

The supervised example used ESM-2 t30 150M embeddings and the representation-complete Core. Five resource-level Core labels were converted into four demonstration classes. `depolymerase`, `endolysin`, and `virion_associated_lytic_enzyme` were retained as separate classes. `multiple_nonexclusive` and `other_phage_envelope_lytic_enzyme` were combined into an `other` demonstration class because the latter contains too few examples for an independent conventional classification example.

The full representation-complete population was split once into 80% training and 20% testing using a stratified split and `random_state=42`. No homology-aware partitioning, sequence-similarity filtering, repeated resampling, cross-validation, or hyperparameter optimization was performed.

Three standard classifiers were evaluated:

- **Logistic Regression**: `StandardScaler`, `class_weight=balanced`, `max_iter=3000`, and `random_state=42`;
- **Linear Support Vector Machine**: `StandardScaler`, `LinearSVC`, `class_weight=balanced`, and `random_state=42`;
- **Random Forest**: 300 trees, `max_features=sqrt`, `class_weight=balanced_subsample`, `random_state=42`, and all available CPU threads.

Headline metrics were accuracy, balanced accuracy, macro-averaged F1, and Matthews correlation coefficient. Class-specific precision, recall, F1, support, confusion matrices, prediction tables, training time, and prediction time were also exported.

Because the split was conventional and not homology-controlled, performance was interpreted strictly as a demonstration that frozen target definitions, stable entity identifiers, released representation matrices, and conventional machine-learning software could be combined reproducibly. The example was not designed to estimate generalization to remotely related phage proteins.

##### S1.5.4 Evidence-aware candidate retrieval

The candidate-retrieval workflow demonstrated rule-based construction of a high-support depolymerase subset using only released fields.

The frozen query configuration was:

- canonical target class equal to `depolymerase`;
- evidence Tier restricted to Tier 1 or Tier 2;
- InterPro annotation required;
- Pfam annotation required;
- existing structure not required;
- no minimum or maximum sequence-length constraint.

Filtering was applied sequentially so that each step could be summarized as a candidate-selection funnel. The sequence of filters was complete Core, canonical depolymerase class, Tier 1–2, InterPro availability, Pfam availability, optional sequence-length constraints, and final selection.

Structural availability was treated as a downstream convenience layer rather than as a biological-quality criterion. Consequently, candidates lacking a PDB or retrievable AlphaFold DB structure were not removed. Representation completeness, UniProtKB mapping, structural availability, and detailed InterPro/Pfam relations were joined after candidate selection for inspection and export.

The workflow produced selected entity tables and FASTA files and allowed structure-ready subsets to be generated separately. No ranking score was calculated and no attempt was made to infer antimicrobial activity, host range, stability, specificity, developability, or experimental desirability. The workflow therefore demonstrates evidence-aware dataset construction rather than predictive candidate prioritization.

### S2 Supplementary Results

#### S2.1 Detailed resource and provenance characterization

##### S2.1.1 Source and observation richness

The final integration layer contained 807,366 source observations originating from seven primary resources. Exact-sequence consolidation yielded 759,105 unique sequence entities, corresponding to 48,261 observation occurrences beyond a one-observation-per-entity representation. The ratio of 1.064 source observations per exact-sequence entity reflects repeated occurrences of identical amino-acid sequences under different database records, accessions, genomes, phages, or source-specific identifiers. These repeated observations were preserved through relation tables and were not discarded during entity consolidation.

The contribution of the individual sources differed substantially in scale and granularity. UniProtKB acquisition produced 227,962 accession-level source observations and 215,075 exact sequences within its source-specific layer. PhaLP represented the largest native source layer, comprising 783,489 protein observations and 751,840 exact sequences represented by those proteins. Genome-centred resources operated over broader protein collections before target-specific candidate extraction; INPHARED contained 265,819 protein observations, from which 5,912 candidate protein observations representing 4,510 exact sequences were retained by its source-specific rules, whereas PhagesDB contained 528,516 annotated proteins and 8,751 direct target candidates representing 4,525 unique candidate sequences. The specialized depolymerase sources were smaller and more focused, comprising 50 positive DePP candidates, 384 DposFinder observations representing 374 exact sequences, and 129 DepoCatalog proteins with 129 unique amino-acid sequences.

These source-layer counts describe the native or candidate-level units produced by each independent acquisition workflow and are therefore not additive. They precede the common multisource observation model and reflect differences in native database structure, source-specific candidate extraction, and within-source exact-sequence redundancy. Their heterogeneity illustrates why PhageLysData retained source observations as a separate analytical layer before defining global exact-sequence entities.

At resource closure, 11,867 entities belonged to the evidence-supported Core, 745,092 to the Prediction Extension, and 2,146 to Context (Supplementary Table **S8**). The Prediction Extension therefore accounted for 98.15% of the integrated exact-sequence space, whereas the Core represented 1.56%. This imbalance was preserved deliberately because the prediction-derived space constitutes a reusable candidate universe but was not treated as equivalent to the non-predictive support required for Core membership.

**Supplementary Table S8**: **Scale and resource-universe composition of PhageLysData v1.0.** Source observations represent standardized source-level occurrences, whereas exact-sequence entities are defined by complete normalized amino-acid identity. Percentages for the three resource universes use the 759,105 exact-sequence entities as denominator.

| Resource component | Count | Fraction |
|---|---|---|
| Source observations | 807,366 | – |
| Unique exact-sequence entities | 759,105 | – |
| Observation occurrences beyond one per entity | 48,261 | – |
| Core | 11,867 | 1.56% |
| Prediction Extension | 745,092 | 98.15% |
| Context | 2,146 | 0.28% |

##### S2.1.2 Cross-source overlap and provenance depth

Exact-sequence redundancy was already evident within several individual source layers before multisource integration. UniProtKB contained 12,887 accession observations beyond its 215,075 exact-sequence entities. PhaLP contained 31,649 protein observations redundant by exact sequence, and its native sequence layer included repeated exact sequences with multiplicities extending beyond ten records. INPHARED candidate extraction produced 5,912 candidate observations but 4,510 exact candidate sequences, while DposFinder contained 384 observations representing 374 exact sequences. The strongest within-source recurrence among the focused candidate layers was observed for PhagesDB, where 8,751 selected proteins represented 4,525 unique exact sequences, reflecting repeated proteins across phage records and annotations.

Global integration retained this multiplicity explicitly. Each exact-sequence entity could therefore remain associated with several source observations, source families, accessions, phages, hosts, and biological assertions. Importantly, observation multiplicity and source-family multiplicity were retained as provenance attributes and were not converted directly into evidence strength. Thus, the occurrence of the same sequence in two databases was distinguishable from two genuinely independent forms of support.

The final public entity tables expose fields such as `source_observation_count`, `source_family_count`, and contributing source-family lists, allowing users to distinguish single-source entities from sequences recovered through several acquisition pathways. The associated provenance relations retain the source and native identifier of every contributing observation. Consequently, exact-sequence consolidation reduces molecular redundancy without removing the multiplicity required to reconstruct how an entity entered the resource.

##### S2.1.3 Evidence-lineage and contextual coverage

The source-specific evidence layers differed not only in scale but also in the number and type of assertions they contributed. UniProtKB query processing generated 494,489 query-derived assertions, which were consolidated into 330,436 independent evidence units after accounting for predefined query-independence groups. PhaLP generated 5,854,678 source assertions and 5,768,924 independent evidence units after consolidation of repeated representations. INPHARED produced 30,412 assertions and 29,560 independent evidence units. The smaller specialized resources generated 440 evidence units for DePP, 3,350 for DposFinder, 863 for DepoCatalog, and 69,882 for PhagesDB.

For sources in which explicit lineage registries were materialized, the resulting evidence was further grouped into 308 lineages for DePP, 755 for DposFinder, 175 for DepoCatalog, and 5,359 for PhagesDB. These values are source-specific and should not be summed as a universal measure of independent biological evidence because each source exposes different native assertion types and dependency structures. Their purpose is to preserve the origin and upstream relationships of individual assertions.

**Supplementary Table S9**: **Scale of source-specific assertion and evidence layers before multi-source evidence integration.** Evidence-unit and lineage definitions are source-specific and are reported to describe provenance depth; counts are not intended as directly comparable measures of biological confidence.

| Source | Assertions | Independent evidence units | Explicit evidence lineages |
|---|---:|---:|---:|
| UniProtKB | 494,489 | 330,436 | – |
| PhaLP | 5,854,678 | 5,768,924 | – |
| INPHARED | 30,412 | 29,560 | – |
| DePP | 440 | 440 | 308 |
| DposFinder | 3,350 | 3,350 | 755 |
| DepoCatalog | 863 | 863 | 175 |
| PhagesDB | 69,882 | 69,882 | 5,359 |

The 2,146 Context entities illustrate the distinction between sequence retention and target assignment. These entities remained part of the resource because they preserve source or biological context but lacked a valid target assertion under the frozen rules. Conversely, Prediction Extension entities retained valid target assignments supported exclusively by prediction-derived information. The coexistence of these three universes therefore preserves contextual and prediction-derived information without forcing either into the evidence-supported Core.

#### S2.2 Detailed Core characterization

##### S2.2.1 Canonical class and evidence-tier composition

The 11,867 Core entities were distributed across four canonical target classes and the derived `multiple_nonexclusive` label. Endolysins dominated the Core, with 9,888 entities (83.3%), followed by 999 `multiple_nonexclusive` entities (8.4%), 712 depolymerases (6.0%), 241 virion-associated lytic enzymes (2.0%), and 27 other phage-envelope lytic enzymes (0.2%).

The evidence-Tier distribution was similarly asymmetric. Tier 3 contained 9,836 entities (82.9%), Tier 4 contained 1,157 (9.7%), Tier 2 contained 784 (6.6%), and Tier 1 contained 90 (0.8%). Independent recomputation of class and Tier counts from the released Core reproduced all nine frozen expected counts exactly, providing a release-facing consistency check for the descriptive characterization.

**Supplementary Table S10**: **Canonical target-class and evidence-Tier composition of the frozen Core.**

| Canonical target class | Entities | Core fraction | Evidence Tier | Entities | Core fraction |
|---|---|---|---|---|---|
| Endolysin | 9,888 | 83.3% | Tier 1 | 90 | 0.8% |
| `multiple_nonexclusive` | 999 | 8.4% | Tier 2 | 784 | 6.6% |
| Depolymerase | 712 | 6.0% | Tier 3 | 9,836 | 82.9% |
| Virion-associated lytic enzyme | 241 | 2.0% | Tier 4 | 1,157 | 9.7% |
| Other phage-envelope lytic enzyme | 27 | 0.2% | | | |

##### S2.2.2 Tier-by-class distributions

The joint Tier-by-class distribution reflected the evidence predicates used to construct the Core (Supplementary Table **S11**). All 90 Tier 1 entities were depolymerases. Tier 2 was approximately balanced between depolymerases and endolysins, containing 383 (48.9%) and 401 (51.1%) entities, respectively.

Tier 3 was dominated by resolved endolysins, which represented 9,474 of its 9,836 entities (96.3%). The remaining Tier 3 entities comprised 220 virion-associated lytic enzymes, 115 depolymerases, and all 27 entities assigned to `other_phage_envelope_lytic_enzyme`. In contrast, Tier 4 was dominated by `multiple_nonexclusive` entities, which accounted for 999 of 1,157 Tier 4 records (86.3%). This pattern is expected from the ontology because entities retaining more than one supported non-predictive target class are not promoted to the single-class Tier 1–3 states.

**Supplementary Table S11**: **Evidence Tier by canonical target-class distribution in the Core.** Values are entity counts; percentages in parentheses represent fractions within each evidence Tier.

| Evidence Tier | Depolymerase | Endolysin | `multiple_nonexclusive` | Other envelope-lytic | Virion-associated lytic |
|---|---|---|---|---|---|
| Tier 1 | 90 (100.0%) | 0 | 0 | 0 | 0 |
| Tier 2 | 383 (48.9%) | 401 (51.1%) | 0 | 0 | 0 |
| Tier 3 | 115 (1.2%) | 9,474 (96.3%) | 0 | 27 (0.3%) | 220 (2.2%) |
| Tier 4 | 124 (10.7%) | 13 (1.1%) | 999 (86.3%) | 0 | 21 (1.8%) |

##### S2.2.3 Sequence-length distributions

Sequence length differed substantially across canonical target classes (Figure **S1**). Depolymerases were the longest group, with a median length of 692 amino acids and a mean of 715.0, compared with a median of 278 amino acids for endolysins. Virion-associated lytic enzymes had a median length of 375 amino acids, while `multiple_nonexclusive` entities had a median of 332 amino acids. The small `other_phage_envelope_lytic_enzyme` group had a median length of 252 amino acids.

Length heterogeneity was substantial within several classes. Endolysin sequences ranged from 19 to 5,007 amino acids, depolymerases from 65 to 4,401, `multiple_nonexclusive` entities from 43 to 2,395, and virion-associated lytic enzymes from 61 to 1,314 amino acids. These distributions explain part of the difference between Core size and the representation-eligible subset because the numerical-representation contract imposed a maximum sequence length of 1,024 residues.

**Supplementary Table S12**: **Sequence-length distributions across Core canonical target classes.**

| Class | $n$ | Mean | SD | Min | Median | Max |
|---|---|---|---|---|---|---|
| Depolymerase | 712 | 715.0 | 346.7 | 65 | 692 | 4,401 |
| Endolysin | 9,888 | 356.6 | 297.0 | 19 | 278 | 5,007 |
| `multiple_nonexclusive` | 999 | 402.1 | 221.4 | 43 | 332 | 2,395 |
| Other phage-envelope lytic enzyme | 27 | 278.0 | 83.1 | 151 | 252 | 448 |
| Virion-associated lytic enzyme | 241 | 454.9 | 214.8 | 61 | 375 | 1,314 |

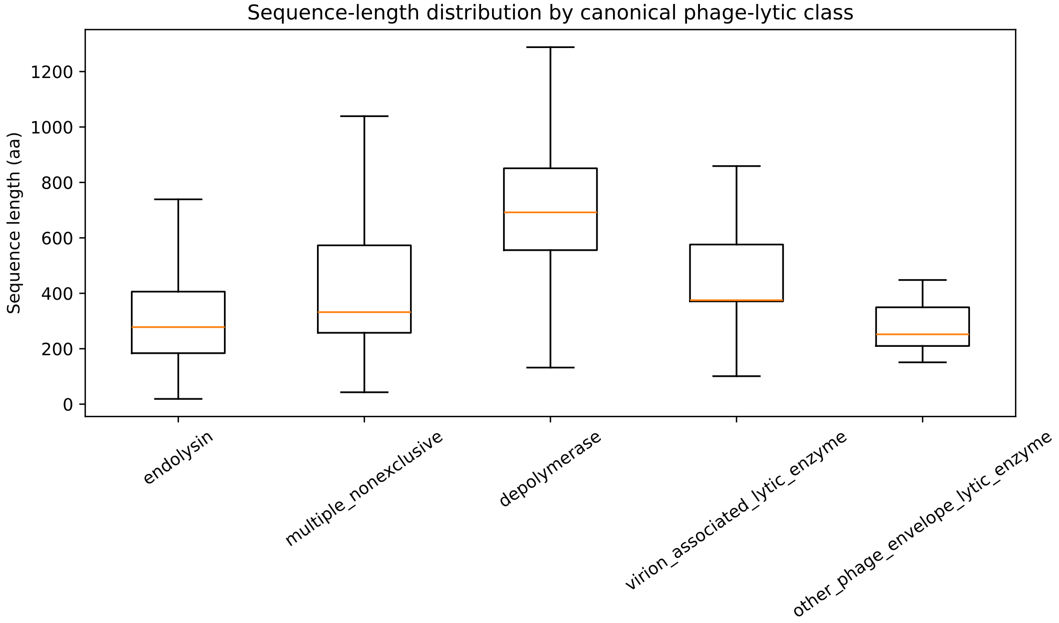

**Supplementary Figure S1**: **Sequence-length distributions across canonical Core classes.** Distribution of amino-acid sequence length for depolymerases, endolysins, `multiple_nonexclusive` entities, other phage-envelope lytic enzymes, and virion-associated lytic enzymes. The distributions are descriptive properties of the frozen Core and were not used to redefine target class or evidence Tier.

##### S2.2.4 Physicochemical-property distributions

The physicochemical layer revealed broad but non-identical property distributions among the canonical target classes (Figure **S2**). Differences in molecular weight closely followed the sequence-length distributions, with depolymerases shifted toward larger molecular weights and endolysins occupying a lower-mass regime.

Net-charge and isoelectric-point distributions also varied among classes. Depolymerases and virion-associated lytic enzymes were shifted toward more negative net charge and lower isoelectric points than the endolysin population. Endolysins showed the broadest distribution toward basic isoelectric points, whereas the medians for depolymerases and virion-associated lytic enzymes remained in the acidic range. The `multiple_nonexclusive` group occupied an intermediate regime.

GRAVY-derived global means were predominantly negative across all groups, indicating that the sequences were not globally dominated by strongly hydrophobic composition. Depolymerases and other phage-envelope lytic enzymes showed less negative median GRAVY values than endolysins and virion-associated lytic enzymes. These distributions describe whole-sequence averages and should not be interpreted as membrane-association or domain-localization predictions.

##### S2.2.5 Annotation and asset coverage by biological class

Annotation and asset availability were not uniform across canonical target classes (Figure **S3**; Supplementary Table **S13**). InterProScan produced at least one match for at least 95.0% of every major class and for all

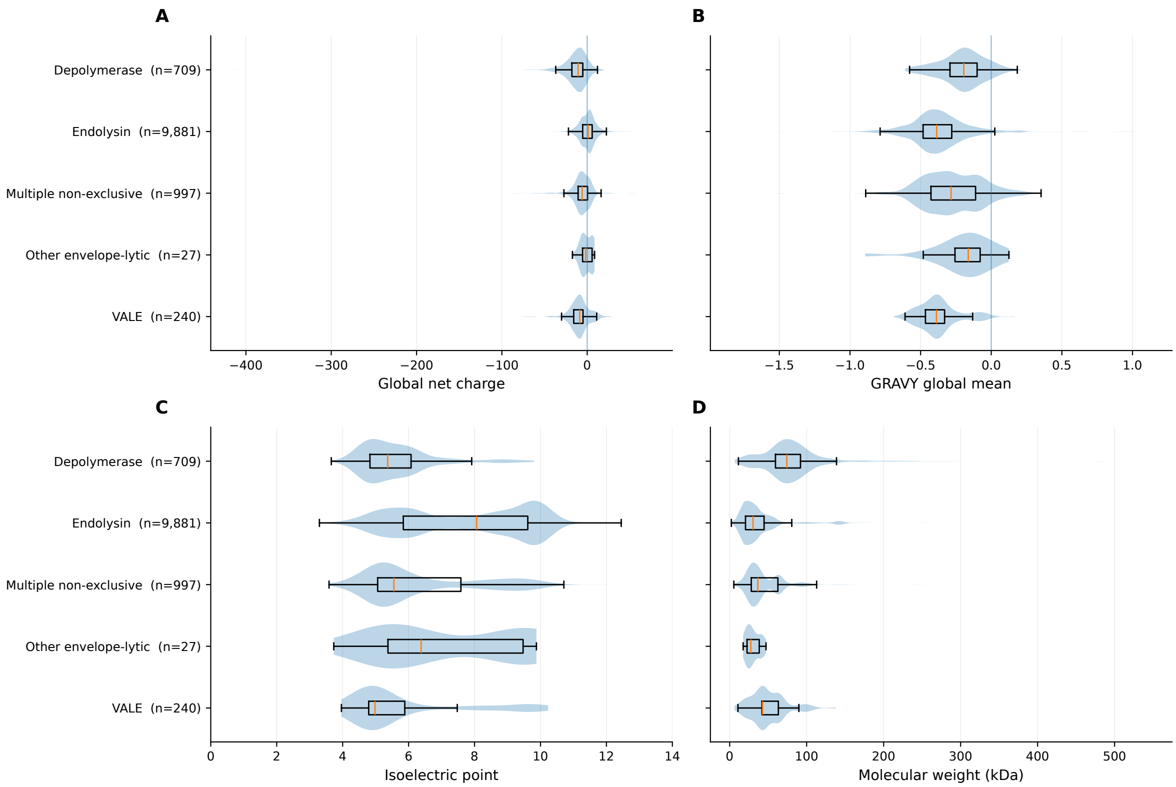

**Supplementary Figure S2**: **Selected physicochemical distributions across Core canonical target classes.** (A) Global net charge, (B) GRAVY full-sequence mean, (C) isoelectric point, and (D) molecular weight. Violin distributions show the observed property distributions, with embedded boxplots summarizing the median and interquartile range; whiskers extend to 1.5 times the interquartile range and individual outliers are not shown. Properties were calculated on the frozen Core without dataset-level normalization or imputation. Sequences incompatible with chemistry-dependent descriptor calculation remain Core entities but are absent from the corresponding property distribution.

27 other phage-envelope lytic enzymes. InterPro assignment was similarly broad, ranging from 86.3% for virion-associated lytic enzymes to 95.1% for endolysins.

Pfam coverage was more variable. It reached 86.3% for virion-associated lytic enzymes and 81.8% for endolysins, compared with 74.3% for depolymerases, 47.0% for `multiple_nonexclusive` entities, and 29.6% for the small other-envelope class. Gene Ontology coverage showed an especially strong class dependence, reaching 63.2% for endolysins but only 8.6% for depolymerases. Pathway mappings were available for 83.0% of endolysins and 85.6% of `multiple_nonexclusive` entities, compared with 67.7% of depolymerases and 40.2% of virion-associated lytic enzymes.

Existing structural coverage remained low for every class, with the highest coverage among endolysins at 7.2%. Numerical representation availability was substantially more complete. Representation completeness ranged from 89.2% for depolymerases to 100% for other phage-envelope lytic enzymes and was identical to representation eligibility within each class, indicating complete materialization for every eligible sequence.

#### S2.3 Functional and structural enrichment

##### S2.3.1 InterProScan annotation landscape

InterProScan processing generated 68,328 raw matches across the 11,867 Core entities. At least one member-database match was obtained for 11,549 entities (97.3%), leaving 318 sequences processed without a match. The match layer represented 15 member databases, 642 unique signatures, and 376 distinct InterPro entries. Pfam contributed 226 unique accessions.

**Supplementary Table S13**: **Annotation and asset coverage by canonical Core class.** Values are percentages of entities within each class. InterProScan match indicates at least one raw InterProScan member-database match; InterPro represents mapped InterPro entries.

| Class | UniProt | IPS match | InterPro | Pfam | GO | Pathway | Structure | Rep. eligible | Rep. complete |
|---|---|---|---|---|---|---|---|---|---|
| Depolymerase | 55.9 | 95.4 | 92.0 | 74.3 | 8.6 | 67.7 | 3.2 | 89.2 | 89.2 |
| Endolysin | 58.7 | 97.5 | 95.1 | 81.8 | 63.2 | 83.0 | 7.2 | 94.8 | 94.8 |
| `multiple_nonexclusive` | 100.0 | 97.9 | 92.7 | 47.0 | 35.1 | 85.6 | 0.4 | 98.9 | 98.9 |
| Other envelope-lytic | 100.0 | 100.0 | 88.9 | 29.6 | 33.3 | 88.9 | 0.0 | 100.0 | 100.0 |
| Virion-associated lytic | 48.1 | 95.0 | 86.3 | 86.3 | 42.7 | 40.2 | 2.5 | 98.8 | 98.8 |

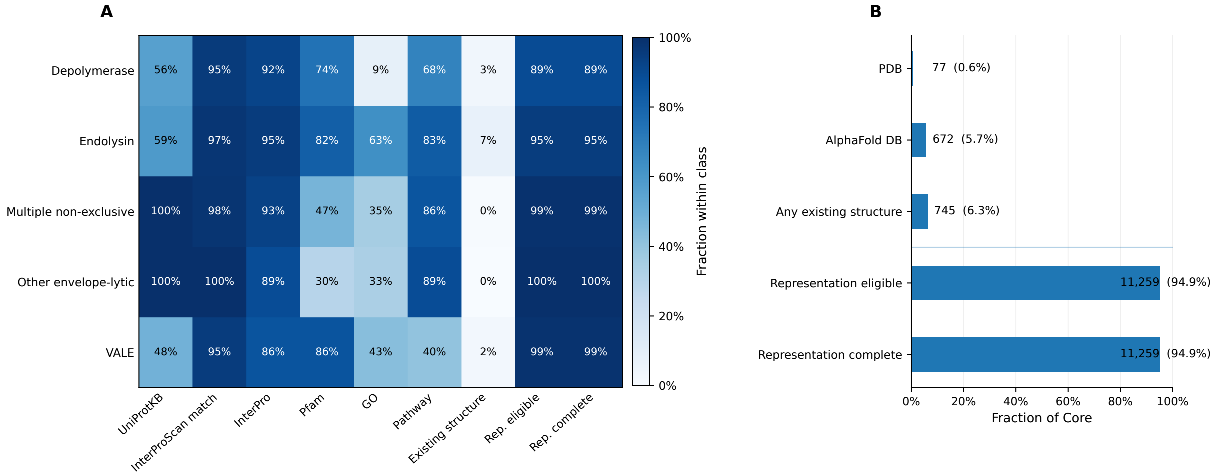

**Supplementary Figure S3**: **Coverage of annotation, structural, and numerical assets in the Core.** (A) Within-class fraction of entities with frozen UniProtKB mapping, at least one InterProScan match, InterPro, Pfam, Gene Ontology, pathway, existing-structure, representation-eligibility, and complete numerical-representation coverage. Values within cells indicate percentages relative to the corresponding frozen Core class. (B) Core-wide availability of mapped PDB structures, AlphaFold DB models, any existing structural asset, representation eligibility, and complete numerical representations. Counts and percentages use the 11,867 frozen Core entities as denominator.

Gene3D provided matches for 10,531 of the 11,549 entities with at least one match (91.2%), followed by SUPERFAMILY for 9,926 (85.9%) and Pfam for 9,300 (80.5%). CDD and PANTHER were represented in 4,788 and 4,542 entities, respectively. These values reflect member-database coverage within the independent InterProScan layer and are separate from corresponding cross-references reported by UniProtKB.

The most frequent InterPro entry was the lysozyme-like domain superfamily (IPR023346), observed in 3,334 entities, corresponding to 29.7% of entities with an InterPro assignment. The N-acetylmuramoyl-L-alanine amidase/PGRP domain superfamily (IPR036505) occurred in 2,095 entities, and the N-acetylmuramoyl-L-alanine amidase domain (IPR002502) in 1,930. Other recurrent entries included the lysozyme domain superfamily, glycoside hydrolase family 24, alpha/beta hydrolase fold, and Peptidase M74/Hedgehog-like zinc-binding domain superfamily.

At the Pfam level, PF01510 (N-acetylmuramoyl-L-alanine amidase) was the most frequent accession, occurring in 1,871 of the 9,300 Pfam-annotated entities (20.1%). PF00959 (phage lysozyme) occurred in 1,536 entities (16.5%). Peptidoglycan-binding, M23 peptidase, soluble lytic transglycosylase, CHAP, glycosyl hydrolase family 25, and LysM domains were also among the recurrent Pfam annotations.

##### S2.3.2 InterPro, Pfam, Gene Ontology, and pathway coverage

The functional landscape differed among canonical target classes. Among the 655 depolymerases carrying an InterPro assignment, the most frequent entries were the pectin lyase fold/virulence-factor superfamily (IPR011050; 341 entities, 52.1%) and pectin lyase fold (IPR012334; 295, 45.0%). Phage-tail-related domains and intramolecular chaperone auto-processing domains were also recurrent. Among Pfam-annotated

**Supplementary Table S14**: **Most frequent independent InterPro and Pfam annotations in the Core.** InterPro percentages use the 11,215 entities with at least one InterPro assignment as denominator; Pfam percentages use the 9,300 entities with at least one Pfam match.

| InterPro | | | | Pfam | | | |
|---|---|---|---|---|---|---|---|
| **ID** | **Description** | *n* | **%** | **ID** | **Description** | *n* | **%** |
| IPR023346 | Lysozyme-like domain superfamily | 3,334 | 29.7 | PF01510 | N-acetylmuramoyl-L-alanine amidase | 1,871 | 20.1 |
| IPR036505 | Amidase/PGRP domain superfamily | 2,095 | 18.7 | PF00959 | Phage lysozyme | 1,536 | 16.5 |
| IPR002502 | N-acetylmuramoyl-L-alanine amidase | 1,930 | 17.2 | PF01471 | Putative peptidoglycan-binding domain | 613 | 6.6 |
| IPR023347 | Lysozyme domain superfamily | 1,613 | 14.4 | PF01551 | Peptidase family M23 | 596 | 6.4 |
| IPR002196 | Glycoside hydrolase family 24 | 1,536 | 13.7 | PF01464 | Transglycosylase SLT domain | 509 | 5.5 |

depolymerases, the most frequent domains included the phage-tail small four-stranded beta-sheet domain (PF27114), endosialidase chaperone (PF13884), and pectate-lyase-like PF12708.

Endolysins showed a different enrichment profile. The lysozyme-like domain superfamily was observed in 3,098 of 9,402 InterPro-annotated endolysins (33.0%), while amidase/PGRP-related entries and glycoside hydrolase family 24 were also frequent. The dominant Pfam assignments were PF01510 and PF00959, followed by peptidoglycan-binding, M23 peptidase, and soluble lytic transglycosylase domains.

The `multiple_nonexclusive` group was enriched for alpha/beta hydrolase fold annotations, which occurred in 559 of 926 InterPro-annotated entities (60.4%), and for phage Gp5-related Pfam domains. Virion-associated lytic enzymes showed strong representation of phage-tail baseplate attachment domains: several Gp16/baseplate-related InterPro entries each occurred in approximately 35% of the InterPro-annotated virion-associated group. The small other-envelope class was dominated by the alpha/beta hydrolase fold, with cutinase-family Pfam support among its Pfam-annotated members. These annotation patterns are descriptive and were not used to redefine the frozen canonical target classes.

**Supplementary Table S15**: **Representative class-enriched InterPro and Pfam annotations.** Percentages are calculated among entities of the indicated class that possess the corresponding annotation type and therefore describe annotation composition rather than whole-class coverage.

| **Class** | **Type** | **Leading annotation** | *n* | **Within-annotated fraction** |
|---|---|---|---|---|
| Depolymerase | InterPro | IPR011050, pectin lyase fold/virulence factor | 341 | 52.1% |
| Depolymerase | Pfam | PF27114, phage-tail small four-stranded beta-sheet domain | 134 | 25.3% |
| Endolysin | InterPro | IPR023346, lysozyme-like domain superfamily | 3,098 | 33.0% |
| Endolysin | Pfam | PF01510, N-acetylmuramoyl-L-alanine amidase | 1,871 | 23.1% |
| `multiple_nonexclusive` | InterPro | IPR029058, alpha/beta hydrolase fold | 559 | 60.4% |
| `multiple_nonexclusive` | Pfam | PF06714, Gp5 N-terminal OB domain | 232 | 49.4% |
| Virion-associated lytic | InterPro | IPR031861, tail baseplate attachment N-terminal barrel | 74 | 35.6% |
| Virion-associated lytic | Pfam | PF16792, tail baseplate attachment N-terminal barrel | 74 | 35.6% |
| Other envelope-lytic | InterPro | IPR029058, alpha/beta hydrolase fold | 24 | 100.0% |
| Other envelope-lytic | Pfam | PF01083, cutinase | 8 | 100.0% |

Gene Ontology mappings were obtained for 6,775 Core entities (57.1%), while pathway mappings were available for 9,662 (81.4%). The most recurrent GO identifiers were associated with peptidoglycan and cell-wall processes and hydrolytic activities, consistent with the dominant endolysin fraction of the Core. Pathway and GO mappings were retained as InterProScan-derived computational annotations and were not interpreted as independent experimental evidence for pathway participation.

##### S2.3.3 Structural coverage from PDB and AlphaFold DB

Existing structural information covered 745 Core entities (6.3%). PDB structures were mapped to 77 entities (0.6% of the Core), while AlphaFold DB models were available for 672 (5.7%). Because four entities had both asset types, the union comprised 745 unique entities.

Structural availability was strongly concentrated in the endolysin class. Endolysins accounted for 712 of the 745 Core entities with an existing structure, corresponding to 7.2% within-class coverage. Depolymerases contributed 23 structure-linked entities (3.2% of the class), virion-associated lytic enzymes six (2.5%), and `multiple_nonexclusive` entities four (0.4%). No existing structural asset was recovered for the 27 other phage-envelope lytic enzymes under the frozen retrieval workflow.

The difference between structural and numerical coverage was substantial. Whereas existing structures were available for only 6.3% of Core entities, 11,259 entities (94.9%) possessed the complete numerical representation layer. Structural availability therefore represents a comparatively sparse complementary layer and was not used as a prerequisite for inclusion or downstream numerical reuse.

#### S2.4 Illustrative reuse analyses

##### S2.4.1 Protein language model latent-space exploration

The released ESM-2 t30 150M embedding contained 11,259 entities and 640 numerical features. PCA retained 50 components, which collectively explained 92.2% of the embedding variance (Figure **S4**A). PC1 explained 22.4% and PC2 9.1%, such that the first two principal components jointly captured 31.5% of the total variance.

The first two PCs already showed non-uniform class organization, although substantial overlap remained (Figure **S4**B). Depolymerases were shifted toward a region distinct from the centre of the endolysin distribution, while virion-associated and `multiple_nonexclusive` entities occupied partially overlapping areas. Endolysins, reflecting their numerical predominance and sequence diversity, occupied the broadest region of the PCA space.

UMAP revealed a more fragmented latent organization with several compact and disconnected regions (Figure **S4**C). Depolymerases formed a conspicuous compact region at low UMAP2 values, whereas endolysins occupied multiple separated structures throughout the projection. Virion-associated lytic enzymes showed both localized regions and overlap with other classes, and `multiple_nonexclusive` entities occurred in several shared regions.

Evidence Tier overlays did not reproduce a simple global ordering of the latent space (Figure **S4**D). The representation-complete population contained 84 Tier 1, 739 Tier 2, 9,309 Tier 3, and 1,127 Tier 4 entities. Tier 3 dominated most regions because it also dominates the underlying Core, whereas Tier 1 and Tier 2 entities were restricted to a smaller number of regions. Tier 4 entities were distributed across several regions occupied by both resolved and multiple-class sequences. These patterns were treated as properties of the selected pretrained representation and projection and not as evidence of homology or experimentally established functional separation.

##### S2.4.2 HDBSCAN clustering of the representation space

HDBSCAN applied to the 50-dimensional PCA-reduced ESM-2 representation identified 53 computational clusters. Of the 11,259 representation-complete entities, 5,688 were assigned to clusters and 5,571 (49.5%) remained classified as noise. Cluster sizes ranged from the minimum imposed by the analysis to 482 entities, with a median cluster size of 86 (Figure **S5**A,B).

Class composition varied considerably among clusters (Figure **S5**C). The largest cluster, C7, contained 482 entities and was 99.2% depolymerase, accounting for a prominent compact region observed in the latent-space visualization. C4 contained 287 entities and was entirely endolysin. Several other large clusters, including C46, C23, C3, C27, C41, C51, and C18, were also composed entirely of endolysins.

Other clusters captured mixed resource labels. C5 contained 264 entities and was dominated by `multiple_nonexclusive` sequences (72.3%), with virion-associated lytic enzymes representing 23.5% and endolysins 4.2%. C12 contained 191 entities and was approximately evenly divided between endolysins (50.8%) and `multiple_nonexclusive` entities (48.7%). Across all non-noise clusters, median canonical-class purity was 1.0 and median normalized class entropy was 0.0, reflecting the large number of class-homogeneous endolysin clusters. These summary values should be interpreted in the context of the strongly imbalanced class distribution.

Evidence-Tier composition followed, but did not exactly reproduce, class composition. C7 contained all four Core Tiers, including 17.4% Tier 1 and 58.1% Tier 2 entities, whereas many pure endolysin clusters were composed almost entirely or exclusively of Tier 3. Clusters enriched for `multiple_nonexclusive` entities were correspondingly enriched for Tier 4.

Annotation coverage also differed markedly across clusters (Figure **S5**D). Several endolysin-rich clusters had nearly complete InterProScan, InterPro, Pfam, GO, pathway, and UniProtKB coverage, while others had complete InterProScan/InterPro support but sparse GO, Pfam, or UniProtKB mapping. Existing structural coverage remained low in most clusters. These patterns demonstrate that the latent representation can be used to identify computationally coherent subsets with distinct annotation profiles, but no cluster-derived functional labels were propagated back into the resource.

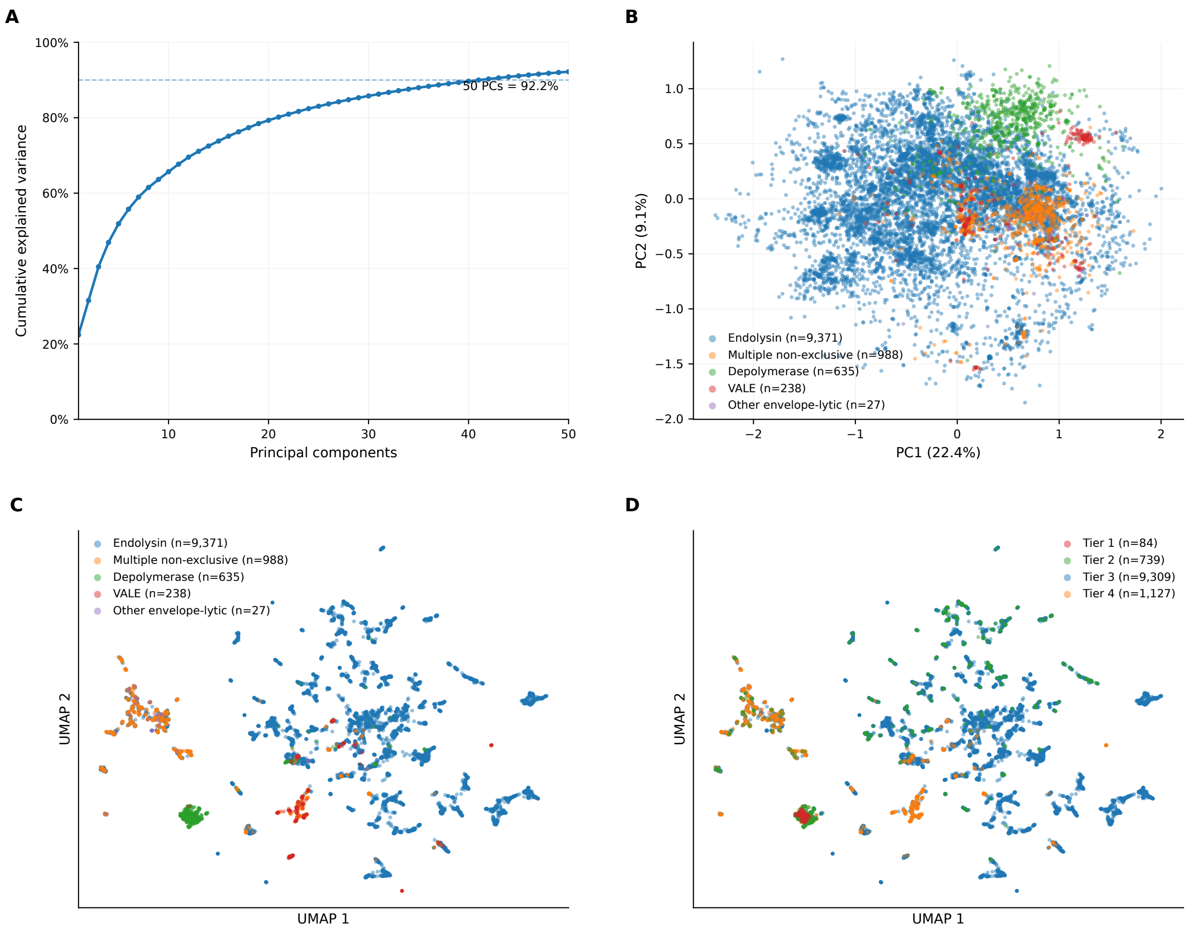

**Supplementary Figure S4**: **Exploration of the released ESM-2 t30 150M latent sequence representation.** (A) Cumulative variance retained by the first 50 principal components. (B) Projection onto the first two principal components, coloured by frozen canonical target class. (C) UMAP visualization generated from the 50-dimensional PCA representation and coloured by canonical target class. (D) The same UMAP coordinates coloured by evidence Tier. Class and Tier metadata were joined after dimensionality reduction and did not influence the projection.

##### S2.4.3 Illustrative supervised classification

The representation-complete Core supported a four-class demonstration task containing 635 depolymerases, 9,371 endolysins, 238 virion-associated lytic enzymes, and 1,015 entities in the combined `other` class. The latter contained `multiple_nonexclusive` and `other_phage_envelope_lytic_enzyme` records. The single stratified split yielded 9,007 training and 2,252 test entities while preserving the strong underlying class imbalance.

All three conventional models recovered substantial class information from the frozen ESM-2 representation (Figure **S6**). Random Forest produced the highest macro-F1 (0.818) and accuracy (0.927). Linear SVM achieved a similar macro-F1 of 0.817 and the highest MCC (0.750), while Logistic Regression produced the highest balanced accuracy (0.877) with macro-F1 0.806 and MCC 0.739 (Supplementary Table **S16**).

Class-specific performance differed more strongly than the aggregate metrics. Depolymerases were recovered with F1 values between 0.950 and 0.980 across the three models, while endolysin F1 ranged from 0.947 to 0.961. Performance was lower for virion-associated lytic enzymes, with F1 values from 0.673 to 0.747, and for the heterogeneous `other` class, with F1 values from 0.615 to 0.645.

The confusion matrices showed that the principal errors involved the smaller and heterogeneous classes. For the Linear SVM, 25.0% of virion-associated lytic enzymes were assigned to `other`, while 16.3% of true `other`

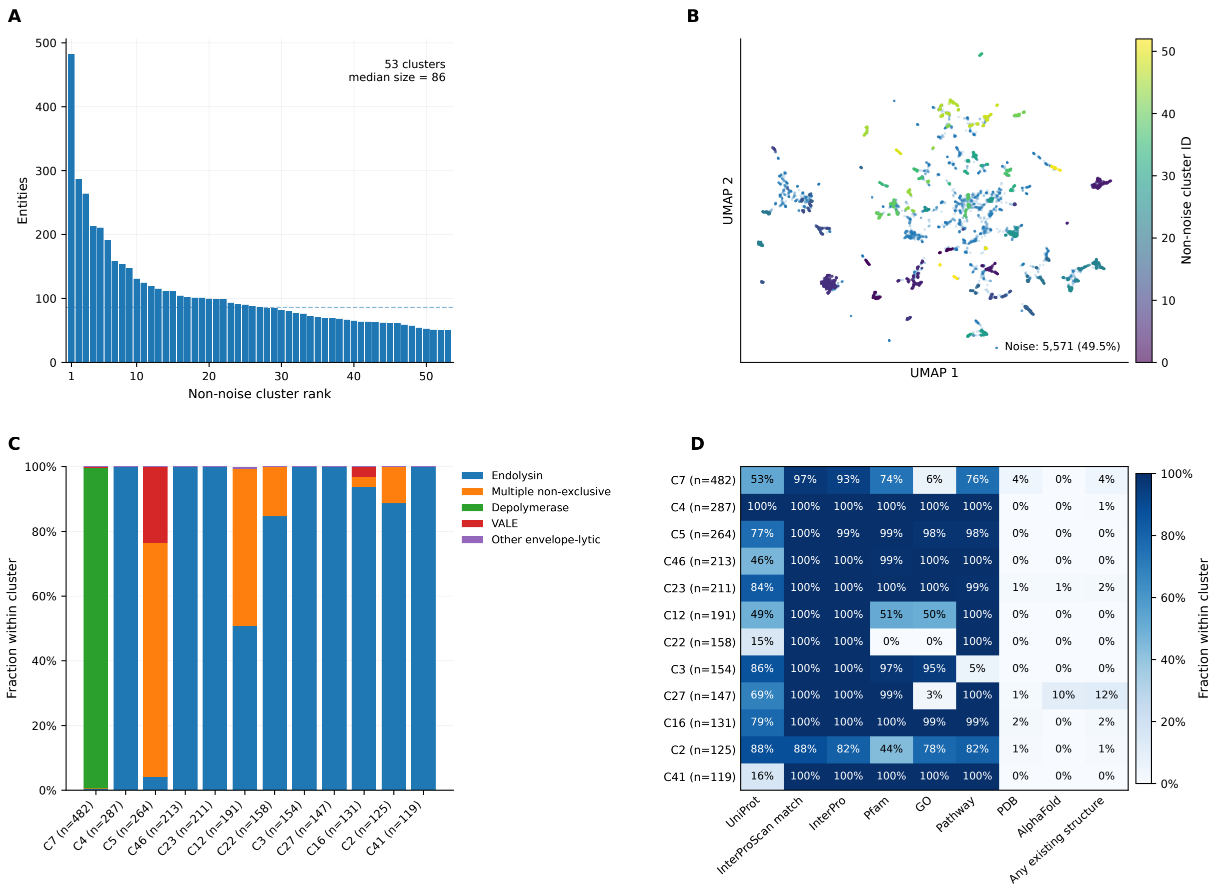

**Supplementary Figure S5**: **Illustrative HDBSCAN grouping of the released ESM-2 representation.** Clustering was performed in the PCA-reduced embedding space and not in UMAP coordinates. (A) Sizes of the 53 non-noise clusters. (B) Cluster assignments visualized using the independently generated UMAP coordinates. (C) Canonical target-class composition of the 12 largest clusters. (D) Annotation and structural coverage across the same largest clusters. Computational clusters were not interpreted as newly validated biological families.

entities were assigned to endolysin. Random Forest recovered 97.4% of endolysins correctly but classified 41.4% of `other` test entities as endolysin. These results demonstrate that the released representations contain information associated with the frozen class labels, while also showing that performance differs markedly across the imbalanced classes.

**Supplementary Table S16**: **Performance of the illustrative four-class classification workflow.** Values are calculated on the single frozen stratified test set. This analysis was not homology controlled and should not be interpreted as a benchmark of remote-sequence generalization.

| Model | Accuracy | Balanced accuracy | Macro-F1 | MCC |
|---|---|---|---|---|
| Random Forest | 0.927 | 0.774 | 0.818 | 0.740 |
| Linear SVM | 0.917 | 0.848 | 0.817 | 0.750 |
| Logistic Regression | 0.903 | 0.877 | 0.806 | 0.739 |

##### S2.4.4 Evidence-aware depolymerase retrieval

The rule-based retrieval workflow started from all 11,867 Core entities and identified 712 canonical depolymerases (Figure **S7**A). Restricting this group to Tier 1 and Tier 2 retained 473 entities, corresponding exactly to the 90 Tier 1 and 383 Tier 2 depolymerases available before functional-annotation filtering.

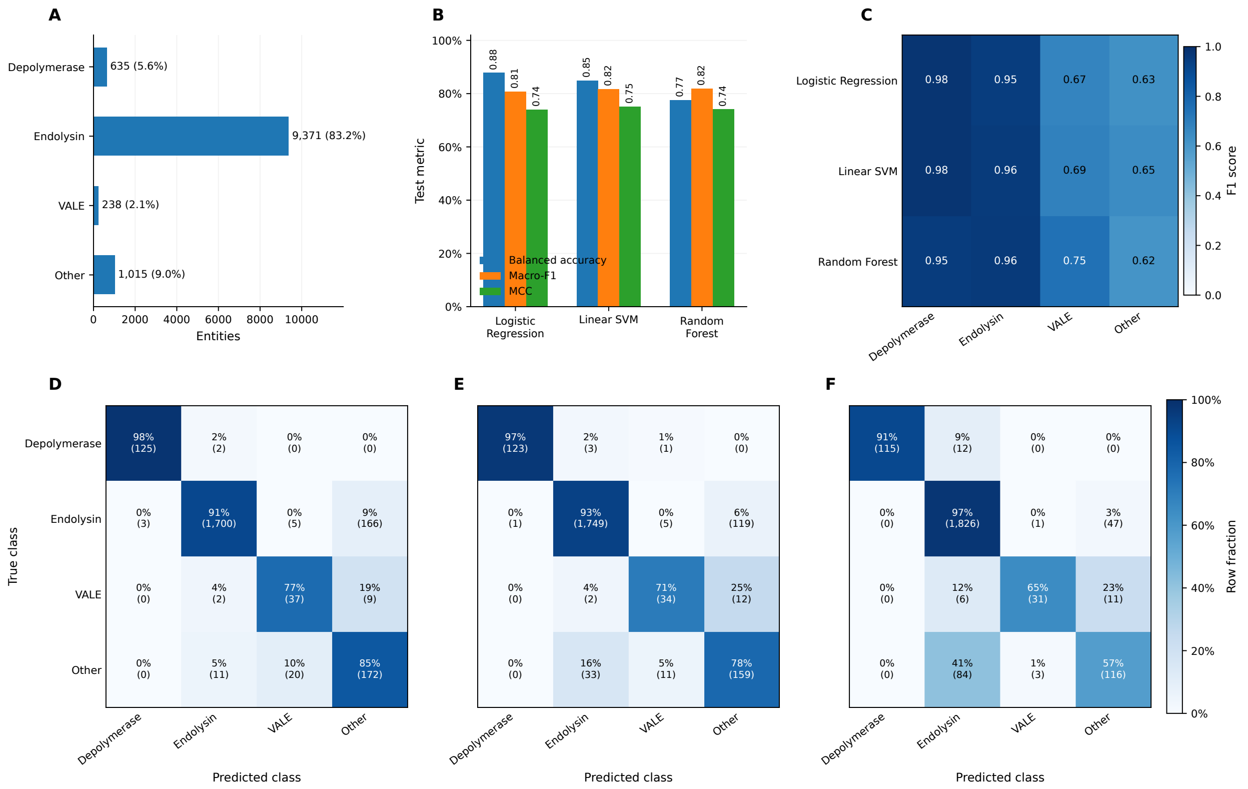

**Supplementary Figure S6**: **Illustrative supervised classification using the released ESM-2 t30 150M representation.** (A) Distribution of the four demonstration classes. (B) Balanced accuracy, macro-F1, and Matthews correlation coefficient for Logistic Regression, Linear SVM, and Random Forest on the frozen test set. (C) Per-class F1 scores for the four demonstration classes. (D–F) Row-normalized confusion matrices for Logistic Regression, Linear SVM, and Random Forest, respectively; each cell reports the row percentage and exact test-set count. The workflow used a single stratified random split without hyperparameter optimization or homology-aware separation.

Requiring an independent InterPro annotation retained 434 entities (91.8% of the Tier 1–2 subset). The additional requirement for Pfam support retained 349 entities, corresponding to 80.4% of the InterPro-filtered set and 49.0% of all canonical Core depolymerases. No sequence-length filter was applied, so all 349 entities passing the evidence and functional-annotation criteria entered the final selected subset. The final set comprised 57 Tier 1 and 292 Tier 2 entities.

The selected proteins had a median sequence length of 772 amino acids and ranged from 175 to 2,417 residues. Their mean source-family count was 1.37 and the maximum was four source families, illustrating that the selected set included both single-source and multisource entities. Chemistry-dependent physicochemical properties were available for 348 of the 349 selected proteins; one sequence contained a non-canonical residue and therefore retained only compatible descriptors.

The asset layer remained heterogeneous after biological selection (Figure **S7**B). By definition, all 349 selected proteins possessed InterPro and Pfam annotations. Complete numerical representations were available for 304 entities (87.1%), while frozen UniProtKB mappings were available for 138 (39.5%). Fourteen selected depolymerases (4.0%) had at least one existing PDB or AlphaFold DB structural asset. The resulting structure-ready FASTA therefore represented a small subset of the evidence-selected population and was distributed separately from the full selected sequence set.

The workflow thus produced an evidence- and annotation-constrained subset without assigning an optimization score. Structural availability, numerical representations, UniProtKB mappings, and physicochemical properties were treated as reusable downstream assets rather than criteria defining biological quality. The selected 349 entities therefore constitute a reproducible example dataset, not a ranked list of preferred experimental candidates.

**Supplementary Table S17: Evidence-aware depolymerase retrieval funnel and final asset coverage.**

| Selection or asset | Entities | Fraction |
|---|---|---|
| All Core entities | 11,867 | 100.0% of Core |
| Canonical depolymerase | 712 | 6.0% of Core |
| Tier 1 or Tier 2 | 473 | 66.4% of depolymerases |
| InterPro available | 434 | 91.8% of previous |
| Pfam available / final selection | 349 | 80.4% of previous |
| Final Tier 1 | 57 | 16.3% of selected |
| Final Tier 2 | 292 | 83.7% of selected |
| Complete numerical representation | 304 | 87.1% of selected |
| Frozen UniProtKB mapping | 138 | 39.5% of selected |
| Existing structure | 14 | 4.0% of selected |

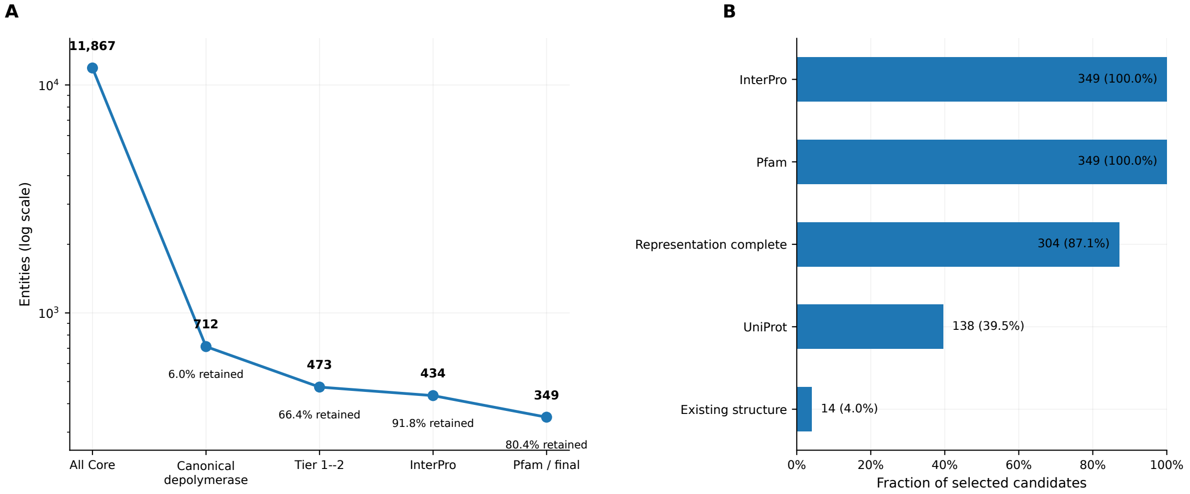

**Supplementary Figure S7: Evidence-aware retrieval of high-support depolymerases from the public Core.** (A) Sequential filtering of the 11,867 Core entities by canonical target class, evidence Tier, independent InterPro annotation, and Pfam support. Entity counts are shown at each stage; percentages after the first stage indicate retention relative to the preceding filter. No sequence-length restriction was applied. (B) Availability of reusable annotations and assets among the 349 selected entities. Existing structural information was not required for selection and was evaluated only after the biological and annotation filters had been applied.

#### References

[1] Uniprot: the universal protein knowledgebase in 2025. *Nucleic acids research*, 53(D1):D609–D617, 2025.

[2] Alexandre Boulay, Victor Németh, Bjorn Criel, Michiel Stock, Bernard De Baets, Clovis Galiez, Elsa Rousseau, Yves Briers, and Roberto Vázquez. Phalp 2.0: extending the community-oriented phage lysin database with a sublyme pipeline for metagenomic discovery. *Database*, 2026:baag033, 2026.

[3] Ryan Cook, Nathan Brown, Tamsin Redgwell, Branko Rihtman, Megan Barnes, Martha Clokie, Dov J Stekel, Jon Hobman, Michael A Jones, and Andrew Millard. Infrastructure for a phage reference database: identification of large-scale biases in the current collection of cultured phage genomes. *Phage*, 2(4):214–223, 2021.

[4] Damian J Magill and Timofey A Skvortsov. Depolymerase predictor (depp): a machine learning tool for the targeted identification of phage depolymerases. *BMC bioinformatics*, 24(1):208, 2023.

[5] Yanxiang Shen, Heyuan Lun, Yumeng Zhang, Zhikang Wang, Cui Tai, Xiaohua Chen, Jiangning Song, Ping He, and Hong-Yu Ou. Dposfinder: an interpretable transformer model for predicting phage-derived polysaccharide depolymerases and their host capsular serotypes. *Genome Medicine*, 2026.

[6] Aleksandra Otwinowska, Sebastian Olejniczak, Agnieszka Latka, Maria Pozniak, Grazyna Majkowska-Skrobek, Barbara Maciejewska, Janusz Koszucki, Vyshakh R Panicker, Sara Jablonska, Mathilde Hulsens, et al. Depocatalog: mapping diversity of 129 recombinantly produced klebsiella phage depolymerases. *Nature Communications*, 2026.

[7] Daniel A Russell and Graham F Hatfull. Phagesdb: the actinobacteriophage database. *Bioinformatics*, 33(5):784–786, 2017.

[8] Gene Ontology Consortium. The gene ontology resource: 20 years and still going strong. *Nucleic acids research*, 47(D1):D330–D338, 2019.

[15] David Medina-Ortiz, Diego Álvarez Saravia, and Julián García-Vinuesa. Sylphy: Protein sequence representation: Encoders, embeddings, and reductions, 2026.
